# IL-15 enhances anti-tumor immunity of sipuleucel-T by activating lymphocyte subsets and reversing immunoresistance

**DOI:** 10.1101/2023.07.23.550213

**Authors:** Muhammad A. Saeed, Bo Peng, Kevin Kim, Ariel Borkowski, Brian Van Tine, Nadeem Sheikh, Tuyen Vu, Daniel LJ Thorek, Todd Fehniger, Russell K. Pachynski

## Abstract

Engineered cell therapies have emerged as a potent therapeutic option for the hematologic malignancies, however solid tumor responses to similar approaches have been modest. The outlier is Sipuleucel-T (sip-T), an FDA-approved autologous cellular immunotherapy for metastatic castration-resistant prostate cancer (mCRPC). To elucidate parameters of the response profile to this therapy, we report the first high dimensional cellular analyses of sip-T using mass cytometry (CyTOF) and show a lymphoid predominance, with CD3+ T cells constituting the highest proportion (median ∼60%) of sip-T, followed by B-cells, and natural killer (NK) and NKT cells.

We hypothesized that treatment of sip-T with homeostatic cytokines known to activate/expand effector lymphocytes could augment efficacy against prostate cancer. Of cytokines tested, IL-15 treated sip-T showed the most significant activation and proliferation of effector lymphocytes, as well as augmentation of tumor cytotoxicity in vitro. Co-culture of sip-T with IL-15 and control or prostate-relevant antigens showed significant activation and expansion of CD8 T and NKT cells in an antigen-specific manner. Adoptive transfer of IL-15 treated sip-T into NSG mice resulted in potent prostate tumor growth inhibition compared with control sip-T. Evaluation of tumor-infiltrating lymphocytes revealed a 2 to 14-fold higher influx of sip-T and a significant increase in interferon (IFN)-γ producing CD8+ T and NKT cells within the tumor microenvironment (TME) in the IL-15 group. In conclusion, we put forward the first evidence that IL-15 treatment can enhance the functional anti-tumor efficacy of sip-T, providing rationale for combining IL-15 or IL-15 agonists with sip-T to treat mCRPC patients.

**Graphical Abstract:** 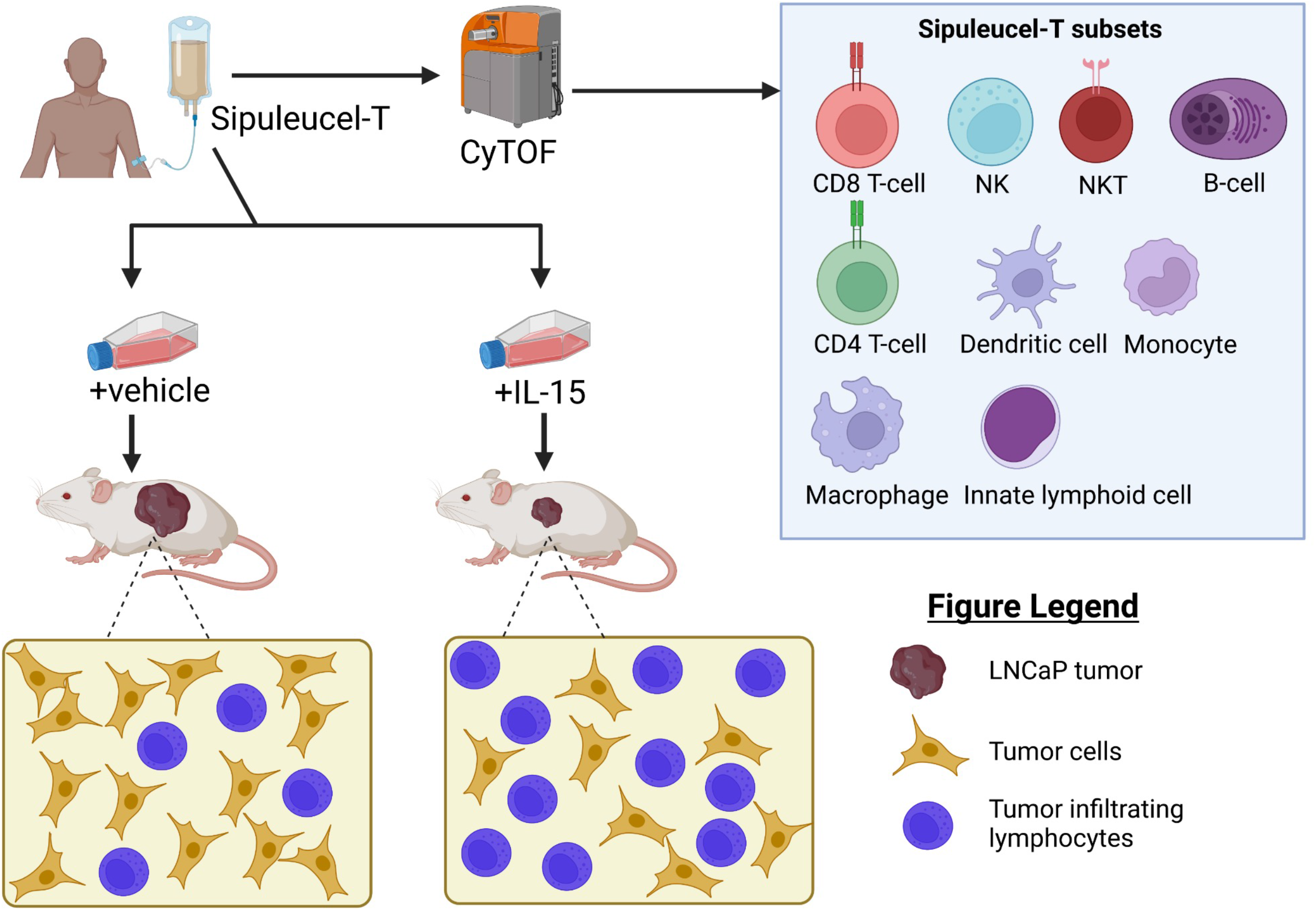

**One sentence summary:** IL-15 treatment can enhance the efficacy/anti-tumor immunity of sipuleucel-T by modulating CD8+ T-cell and CD56+ NKT subsets.

## INTRODUCTION

Prostate cancer is the second most frequently diagnosed cancer among men globally and the second leading cause of cancer-related mortality in the USA, with an estimated 2.68 million new cancer cases and 34500 deaths reported in 2022 (*1*). Immunotherapy represents a promising therapeutic approach for a range of tumor types; however, it may fail to effectively treat prostate cancer (*2*). Sipuleucel-T or sip-T (PROVENGE; Dendreon) is the first and only autologous cellular immunotherapy for use in men with metastatic castration-resistant prostate cancer (mCRPC) approved by the U.S. Food and Drug Administration (FDA) (*3, 4*). In the pivotal randomized, Phase III IMPACT trial (NCT00065442), sip-T prolonged median survival of mCRPC patients by 4.1 months compared with those treated with placebo (*4*). Sip-T is produced by culturing patients’ peripheral blood mononuclear cells (PBMC) ex vivo with a fusion protein PA2024, which combines human recombinant prostatic acid phosphatase (PAP; a commonly expressed prostate tumor antigen) and granulocyte-macrophage colony-stimulating factor (GM-CSF) (*3*). The activated cellular product is then adoptively transferred back into the same patient, and the cycle is generally repeated for two further doses with approximately two-week intervals with freshly made product each time (*2, 3*). This results in enhanced antigen-presenting cell activity and stimulation of cytotoxic T-lymphocytes which lead to increased cellular and humoral responses to PAP as well as generation of secondary immune response to additional tumor antigens (*2, 5, 6*). Sip-T mediated augmentation of the immune response may result in increased clinical benefit observed in patients with mCRPC (*2, 4–6*). Despite its proven therapeutic activity as a single agent, sip-T fails to induce clinically demonstrable responses (i.e., prostate specific antigen (PSA) or radiographic responses) in the majority of mCRPC patients (*2*). Thus, strategies to improve sip-T efficacy in mCRPC patients are needed.

Cytokines can stimulate the activity, survival and proliferation of NK and T cells that mediate immune response against tumors (*7*). Interleukin-15 (IL-15) belongs to the γ-chain family of cytokines, which also includes IL-2, IL-4, IL-7, IL-9 and IL-21 (*8*). These endogenous homeostatic cytokines can stimulate the proliferation and activation of lymphocytes, and augment lymphocyte migration from blood into lymph nodes and peripheral organs (*9, 10*). IL-15 receptor (IL-15R) shares the same β and γ chains with IL-2. However, unlike other secretory cytokines, IL-15 primarily exists bound to IL-15Rα (typically on the surface of myeloid cells), which is trans-presented to the βγ-receptor-positive responding NK or CD8+ T cells (*8, 11*). Therefore, IL-15 is considered crucial for the development, function and homeostasis of CD8+ T cells, NK cells, and NKT cells (*11*). Furthermore, IL-15 exhibits superior activity over IL-2 in terms of weaker Treg-stimulation, lower activation-induced cell death (AICD), lower vascular endothelial toxicity, and stronger expansion of NK and CD8+ T cells (*12*). Within tumor tissues, IL-15 plays a crucial role in establishing a regular number of CD8+ T cells and NK cells (*11, 12*). Low expression levels of IL-15 within tumor microenvironment (TME) have been associated with decreased T cell proliferation, higher tumor recurrence, decreased patient survival and poor prognosis (*13, 14*). Given the ability of human recombinant IL-15(rhIL-15) to expand and enhance lymphocyte populations and their functions, we hypothesized that the treatment of sip-T with rhIL-15 would result in improved anti-tumor efficacy of sip-T. This motivated the current undertaking of the first high dimensional analysis of sip-T composition using mass cytometry (CyTOF) and report of pre-clinical data delineating the effects of IL-15 on sip-T phenotype and efficacy, both in vitro and in vivo. We observed a significant activation and expansion of sip-T with a significant improvement in tumor cytotoxicity following IL-15 treatment in vitro. The adoptive transfer of IL-15 treated sip-T into NSG mice bearing subcutaneously implanted LNCaP human prostate cancer xenografts significantly suppressed tumor growth compared with those treated with control sip-T. Importantly, reduced tumor growth by IL-15 treated sip-T was accompanied by increased influx of sip-T as well as expansion/activation of CD8+ T cell and CD56+ NKT within the tumor microenvironment. RNAseq analyses of the TME revealed upregulation of antigen presentation, lymphocyte activation, and lymphocyte mediated immunity pathways in the IL-15 treatment group; qPCR confirmed increased expression of several key genes involved in anti-tumor immunity (e.g., *IFNG*, *GZMB*, *LAMP1* etc.). Taken together, these data show the addition of IL-15 to sip-T therapy significantly improves its anti-tumor efficacy in preclinical modeling, and provides the rationale for exploring this combination in a human clinical trial in patients with mCRPC.

## RESULTS

### Patient demographics

Sipuleucel-T (sip-T) from 11 metastatic castrate-resistant prostate cancer (mCRPC) patients were included in this study for CyTOF analysis. 14 sip-T samples analyzed by CyTOF consisted of first (n=6), second (n=3), and third (n-5) dose groups. The demographics and related information for these patients are summarized in Table 1. Most patients (n=10, 91%) were of Non-Hispanic Caucasian background while one (n=1, 9%) was Non-Hispanic Black. The age range was 60-82 with a median age of 69 years. All patients except one had confirmed mCRPC and had been treated for metastatic hormone-sensitive prostate cancer (mHSPC) with at least androgen deprivation therapy (ADT), which was maintained during treatment with sip-T. One patient had localized prostate cancer and no prior ADT, but was treated on a clinical trial with sip-T. Patients’ prostate specific antigen (PSA) levels immediately prior to treatment with sip-T ranged between 0.14-57.47 with median of 2.89. Median alkaline phosphatase levels were 83 U/L (range 51-180; normal values 40-130) and hemoglobin 12.6 g/dL (range 9.5-17; normal values 13-17.5). For 73% (n=8) of the patients, sip-T was the first treatment they received for mCRPC disease. Overall, these data indicate that the patient cohort had less heavily pre-treated mCRPC disease and generally had good prognostic features as evidenced by laboratory values and performance status (*15–17*).

### Phenotypic profiling of sipuleucel-T by mass cytometry (CyTOF)

To date, there has been limited data using mass cytometry (CyTOF, cytometry by time of flight) to characterize peripheral blood from patients treated with sip-T (*2, 18*). To our knowledge, there has been no CyTOF evaluation of the sip-T product itself, prior to infusion into patients. Thus, we first performed high dimensional analysis of sip-T to phenotype the cellular contents of this therapy. We designed a comprehensive CyTOF panel that included markers for the identification of major immune cell types, in addition to markers that define their subgrouping and activation/exhaustion states (table S1). A total of 14 sip-T samples from 11 individual mCRPC patients were processed for CyTOF acquisition (Fig. 1, S1). Our results show that majority of sip-T (> 99%) were non-granulocytic leukocytes (CD45+ CD66b-), and only a small proportion of granulocytes (CD45+CD66b+) with an average of less than 0.1% of total sip-T. To spatially visualize the immune cell profiles of sip-T, a UMAP spatial heatmap analysis was applied to CD45 + cells (Fig. 1A). This neighborhood embedding technique allowed us to visualize groupings of cells based on the expression of all major markers in sip-T. CyTOF analysis revealed that CD3+ T cells constituted the highest proportion (median, range: 63%, 9-89%) of sip-T, followed by B-cells (4%, 1-82%), natural killer (NK) cells (4%, 1-18%), NKT (2%, 1-7%) and monocytes (1%, 1-37%), as well as a small proportion (<1%) of conventional dendritic cells (cDC), innate lymphoid cells (ILC), plasmacytoid DC (pDC), and myeloid-derived suppressor cells (MDSC). The majority of CD8+ T cells (Fig. 1D) were either CD45RA+CCR7-effector (median, range: 43%, 9-70%), CD45RA-CCR7-effector memory (EM; 23%, 11-59%) and CD45RA+CCR7+ naive (22%, 4-50%), while most CD4+ T cells (Fig. 1C) belonged to CD45RA-CCR7+ central memory (median, range: 45%, 24-78%), EM (27%, 6-61%), and naive (22%, 1-50%) subtypes. Other notable CD4+ subpopulations included Th1 (Tbet+), Th2 (GATA3+), Th17 (RORγt+) and regulatory T cells (Tregs; CD127-CD25+FoxP3+). Interestingly, CD25-FoxP3+ Tregs constituted a higher proportion (median, range: 7%, 2-42%) of sip-T than classical CD25+FoxP3+ Tregs (median, range: 2%, 1-21%). The majority of monocytes (median, range: 86%, 78-97%) were CD86+, suggesting a potential role in inducing lymphocyte activation (Fig. 1F) (*19*). Likewise, the majority (87%, 27-92%) of these cells were positive for the macrophage marker CD68 (*20*). Variable expression of immune checkpoint molecules, including PD1, PD-L1, CTLA4, VISTA, ICOS and TIM3, were found among different sip-T subsets (fig. S1). Of note, within the lymphoid component, CD4 T cells and ILCs had the highest percentages of cells positive for PD1 expression, while cDCs showed the highest within the myeloid populations. Interestingly, PD1 expression on DCs has been shown to suppress CD8 T cell anti-tumor activity (*21*). PD-L1 expression was highest in myeloid cells, with approximately 40% of monocyte populations expressing detectable levels of PD-L1. CTLA4 and VISTA expression was generally low (<1%) on CD4 and CD8 T cells, whereas ICOS expression on CD4 T cells varied and in some samples was above 20% of the population. Interestingly, a relatively large percentage of CD8 T cells showed expression of the immunoregulatory marker TIM3, with a median of ∼22% and ranging up to ∼75% of the population. TIM3 expression has been shown on PSA-specific CD8 T cells, indicating an exhausted phenotype (*22*). Overall, our CyTOF data shows for the first time detailed phenotypic leukocyte subsets of sip-T, and identify potential therapeutic immune checkpoint targets within sip-T.

**Fig. 1.**
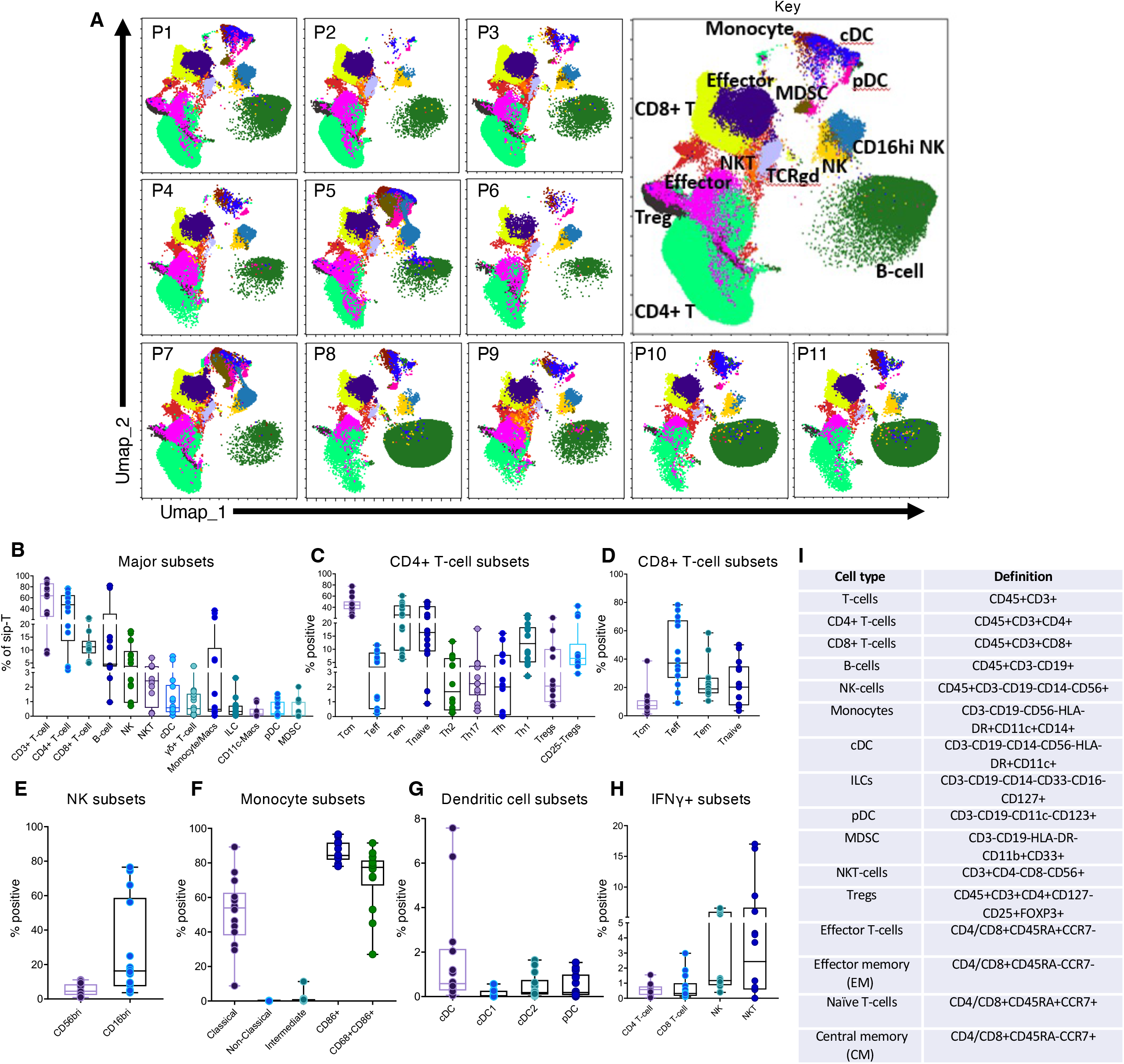
High throughput (CyTOF) profiling of sipuleucel-T from mCRPC patients. Fourteen sip-T samples from 11 individual metastatic castrate-resistant prostate cancer (mCRPC) patients were analyzed by mass cytometry (CyTOF). **(A)** UMAP dimensionality reduction plots of CyTOF data of major immune cell subsets of sip-T from individual patients (p, patient). Annotation key shows major cell types in sip-T represented by different colors. **(B)** Compositional diversity of major immune cell populations of sip-T represented by boxplots. Each patient sample is denoted by an individual symbol. **(C)** CD4+ T-cell subsets. Tcm, central memory T-cells; Teff: effector T-cells; Tem, effector memory T-cells; Tfh, follicular helper T-cells; Tregs, regulatory T-cells. **(D)** CD8+ T-cell subsets **(E)** Natural killer (NK) cell subsets **(F)** Monocyte/macrophage subsets. **(G)** Dendritic cell (DC) subsets. cDC, conventional dendritic cell; pDC, plasmacytoid dendritic cell. **(H)** IFNγ+ cells among various sip-T subsets. **(I)** Definition of various immune subsets of sip-T analyzed in this study.

### IL-15 expands and activates sipuleucel-T lymphocyte populations, enhancing anti-tumor efficacy

We next sought to evaluate whether lymphocyte homeostatic cytokines (IL-7, IL-15, and IL-21) could enhance sip-T efficacy, as they are known to stimulate the proliferation and activation of T and NK cells, and augment T cell migration from blood into lymph nodes and peripheral organs (*9, 10*). While recombinant human IL-7 has been tested in a clinical trial with sip-T and shown to augment immune responses (*2*), no other cytokines have been studied in combination with sip-T either clinically or pre-clinically. To assess the effect of cytokines on sip-T phenotype, a 7-day stimulation of sip-T (n=3 individual patients) was performed using 50 ng/ml of each of IL-7, IL-15, and IL-21 followed by analysis with CyTOF (Fig. 2, A-E). IL-15 treated sip-T showed pronounced expansions of CD8+ T cell, CD56+ NKT and NK cell populations which were significantly higher (up to ∼2-fold) than those of IL-7 and IL-21 treated sip-T (Fig. 2, A-B). Furthermore, IL-15-treated sip-T exhibited a pronounced increase in IFN-γ+, CD107+ (*LAMP1*, a marker of degranulation), and Ki67+ (proliferation) cells in CD8+ T cell, NKT and NK populations (Fig. 2, C-E), with significant increases seen in most of these markers compared to the IL-7 and IL-21 groups (Fig. 2, C-E).

**Fig. 2.**
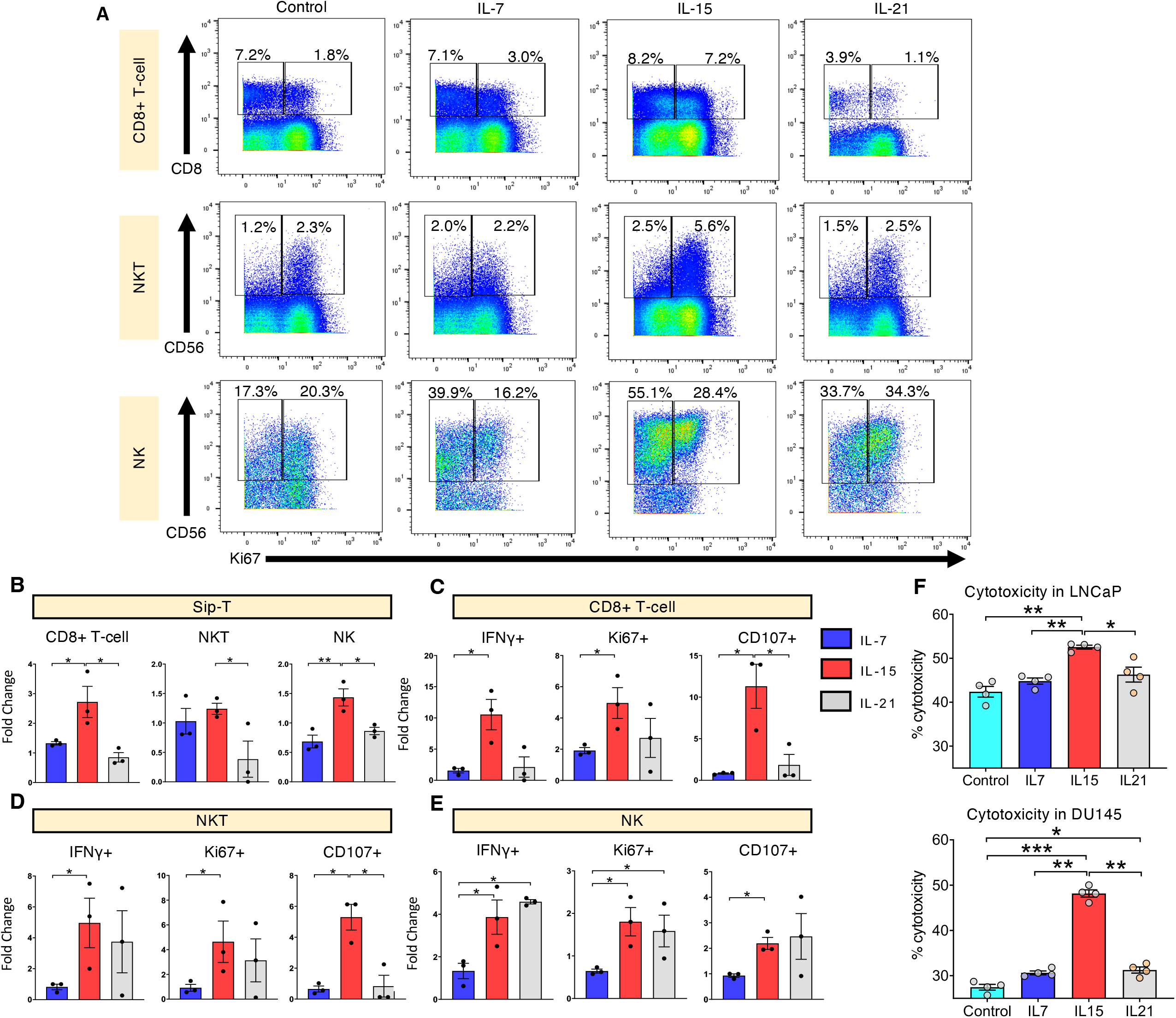
IL-15 exhibits superior activity in enhancing sipuleucel-T efficacy *in vitro*. Sip-T from individual mCRPC patients were treated with 50 ng/ml each of IL-7, IL-15, IL-21 or media only control on day 0, 2, 4 and 6 and analyzed at day 7. **(A)** Representative dot plots of CyTOF-based quantification of cytokine treated sip-T (n=3). Only major immune cell subsets (i.e., CD8+ T-cell, CD56+NKT and NK cell) affected by cytokine-treatment are shown. **(B-E)** Bar graphs show fold change, relative to control, in the composition of sip-T following cytokine treatment. IL-15 treated sip-T showed significantly higher expansion and activation of CD8+ T-cell, NKT and NK populations compared to those of IL-7, IL-21, or both (mean ± SEM shown). **(F)** Cytotoxicity assays. Cytokine treated sip-T were washed and co-cultured for 24 hours with CFSE-labeled LNCaP or DU145 (4:1) and analyzed by flow cytometry. The data represents one of the three independent experiments (mean ± SEM shown). IL-15-treated sip-T exhibited significantly higher cytotoxicity in both tumor lines tested compared with controls or other cytokines tested. *, *P < 0.05*; **, *P < 0.01*; ***, *P < 0.001* by 1-way ANOVA with Tukey’s post hoc test.

To test whether cytokine treatment had a functional impact on sip-T efficacy, we performed cytotoxicity assays using two common prostate tumor cell lines LNCaP (prostate specific membrane antigen, PSMA positive) and DU145 (PSMA negative). Although IL-21 treatment enhanced sip-T mediated cytotoxicity in DU145 compared to control, IL-15 treated sip-T showed significantly higher cytotoxicity in both tumor lines compared to control, IL-7, and IL-21 (Fig. 2F). We then assessed a real-time visualization of the effect of IL-15 on sip-T cytotoxicity, using the IncuCyte® imaging assay. IL-15 treated sip-T showed significantly higher tumor cytotoxicity compared with the patient-matched unstimulated controls over time in both LNCaP and DU145 cell lines (fig. S2). Overall, these data indicate that IL-15 is more effective than IL-7 and IL-21 in expansion and activation of sip-T effector lymphocyte subsets, as well as enhancing its cytotoxic activity against prostate cancer.

### IL-15 treatment enhances proinflammatory response and anti-tumor signatures of sip-T

To further identify mechanisms of IL-15-enhanced efficacy of sip-T, gene expression profiles were analyzed by whole transcriptome RNA sequencing (RNA-seq). Sip-T (n=5 patient samples) was treated with IL-15 or vehicle and analyzed at day 7 post the start of treatment. RNA-seq data showed a differential upregulation of a larger set of genes (504 genes in total) in IL-15 treated sip-T compared with only 307 genes in the control sip-T group (Fig. 3A). 12569 genes were equally regulated between IL-15 treated and control groups. The functional characterization of the differentially expressed genes was performed using GO term enrichment analysis. IL-15 activated and repressed DEGs were separately classified for the GO category “biological process”. IL-15 treatment resulted in enrichment of signatures pathways involved in lymphocyte activation, proliferation, cytotoxicity, downstream antigen receptor signaling, lysosome/autophagosome, secretory/zymogen granules, and antigen and cytokine binding (Fig. 3B), in line with our previous data showing IL-15-mediated activation of T and NK/T cells. We then identified 28 genes relevant to lymphocyte activation/pro-inflammatory response that were common within the upregulated pathways (Fig. 3B), and compared them between control and IL-15 treatment in a heat map (Fig. 3C). Detailed information for significantly upregulated genes and enriched pathways is provided in table S2. We saw significant upregulation of genes involved in effector T cell and NK/T cell development, differentiation, proliferation and activation (*CD3d*, *CD3e*, *CD8a*, *IL2RA*, *IL2RB*, *IRF1*, *IRF5*, *NKG7*, *KIR3DL2*, *SLAMF1*, *ZAP70*, *TNFA*, *EOMES*), lymphocyte trafficking (*CCL3*, *CCL5*), antigen recognition, binding, processing and presentation (*HLA*-B, *HLA*-*DRB1*, *AIM2*, *NLRP3*, *NLRC3*, *RIPK3*, *CD247*) and anti-tumor /cytotoxic/phagocytic activity (*FASLG*, *GZMA*, *GZMB*, *GNLY*, *IFNG*, *TNFSF10*). To confirm whether IL-15-mediated effects on the sip-T transcriptome were translated at the protein level, we performed a Legendplex ELISA to measure secreted proteins from differentially expressed genes in the RNA-seq data. After a 7-day stimulation with IL-15, cell culture supernatants were tested for a variety of proinflammatory and regulatory cytokines and proteins (Fig. 3D). With IL-15 treatment of sip-T, the highest increase (fold change compared with untreated sip-T control) was observed in IL-2 levels (mean±SEM; 15.9±5.4) followed by IFN-γ (2.8±0.8), sFAS-L (2.6±0.4), sFAS (2.0±0.2), granzyme B (2.5±0.4), perforin (1.8±0.2), TNFα (1.7±0.3), granzyme A (1.7±0.2), and granlulysin (1.3±0.1). Interestingly, levels of the immunosuppressive cytokine IL-10 were significantly decreased by IL-15 treatment. No significant changes were observed in IL-17A and IL-6 levels between the two groups. Together, these data indicate that IL-15 favorably modulates sip-T by significantly upregulating several key pro-inflammatory genes and pathways, with resultant increases in secreted anti-tumor effector proteins/cytokines.

**Fig. 3.**
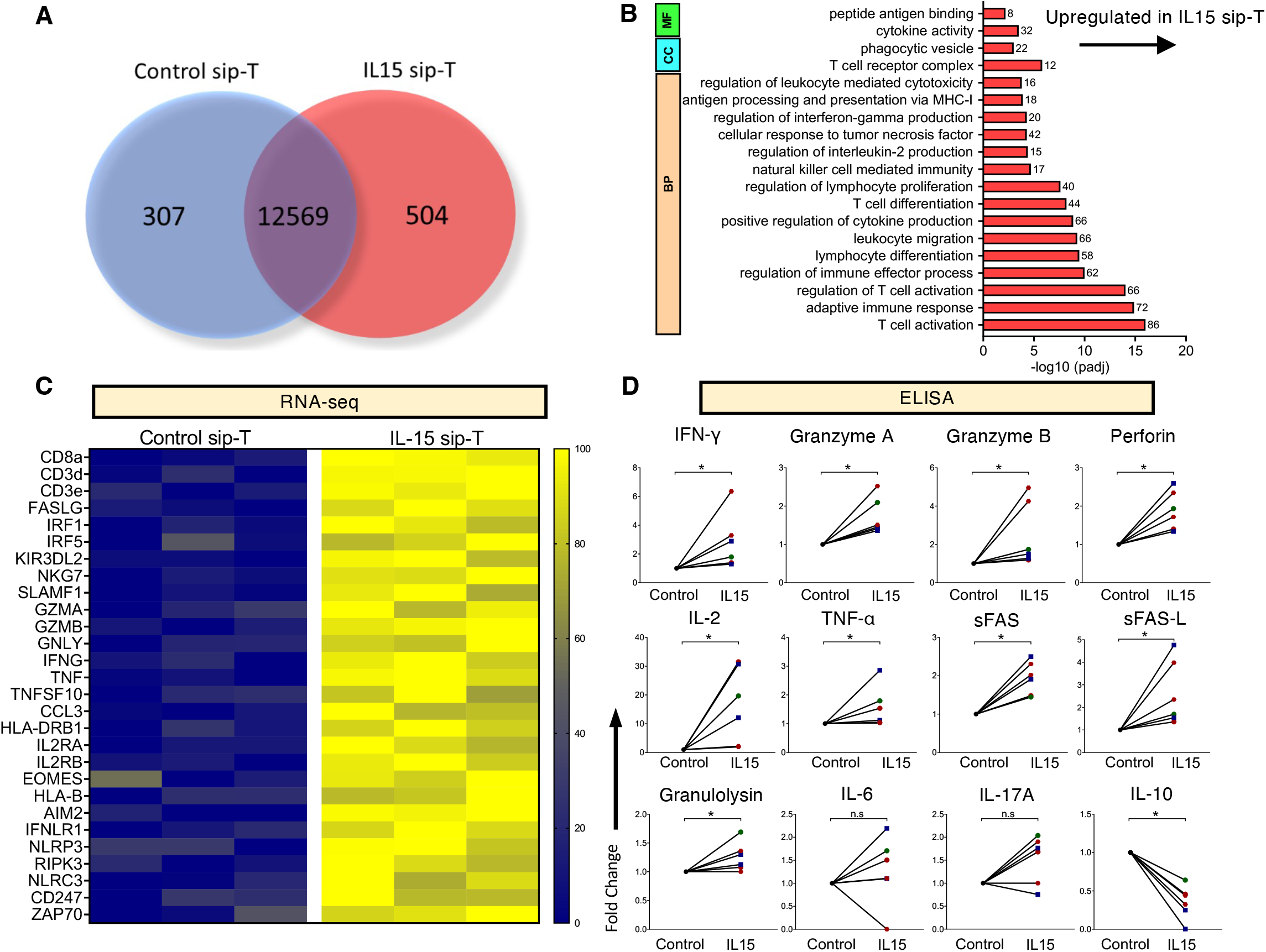
Functional analysis of IL-15 treated sip-T. Sip-T from individual mCRPC patients were treated with IL-15 (50 ng/ml) or control (media only) on day 0, 2, 4, 6 and then harvested at day 7. (A-C) Bulk RNA-seq of sip-T (n=5 patients). **(A)** Venn diagram of differentially expressed genes between IL-15 treated and control sip-T. **(B)** Differential gene pathway analysis of RNA-seq between IL-15 treated and control sip-T. IL-15 treated sip-T showed a significant enrichment in pathways involved in lymphocyte activation, proliferation, lymphocyte-mediated cytotoxicity, downstream antigen receptor signaling, lysosome/autophagosome, secretory/zymogen granules, and antigen and cytokine binding. **(C)** Heatmap displaying differential gene expression of activation/effector markers also showed that many of these genes were upregulated in IL-15 treated sip-T. **(D)** ELISA-based estimation of cytokine secretion from IL-15 treated sip-T (n = 6 from 3 patients). Significantly elevated levels of pro-inflammatory cytokines (IFN-γ, granzyme A/B, perforin, IL-2, TNF-α, sFAS, sFAS-L, and granulysin) and reduced levels of IL-10 were exhibited by IL-15 treated sip-T compared with controls.

### IL-15 expands tumor specific cytotoxic T cell subsets of sip-T

We next wanted to determine if treatment with IL-15 could enhance responses of tumor-antigen specific lymphocyte effector populations. To test this, we co-cultured sip-T with PA2024, which contains the tumor antigen PAP, or control peptide (CEFT) alone or in combination with IL-15. We looked at cell expansion and proliferation (by Ki67), as well as activation (by IFN-γ and CD107) by FACS. Exposure to control peptide or PA2024 alone did not significantly expand CD8+ T or CD56+ NKT cells, however the addition of IL-15 expanded these populations ∼1.8 fold (Fig. 4 A, B). There was no significant difference in proliferation (total cell number or Ki67) or activation (IFN-γ or CD107) in IL-15 treatment groups with the addition of control peptides compared to IL-15 alone. Importantly, we saw significant increases in proliferation and activation of both CD8+ T and NKT cells in the presence of specific tumor antigen (PA2024), compared to IL-15 with or without control peptides. Interestingly, only the combination of IL-15 and PA2024 resulted in a significant increase in CD8+ T cell CD107 expression (Fig. 4 A, B). These data suggest that IL-15 not only non-specifically expands and activates CD8+ T and NKT cell sip-T populations, but acts to augment tumor antigen specific responses in these populations. In NKT cells, IFN-γ and CD107 expression was significantly increased only with the combination of IL-15 and PA2024 (Figure 4B), suggesting potential synergistic effects. No significant differences were observed in CD4+ T cell subset frequencies following PA2024 treatment in the presence of IL-15 (data not shown). Interestingly, IL-15 significantly expanded PA2024-specfic central memory and effector memory subsets of CD8+ T cells (Fig. 4A) but not effector or naïve subsets (data not shown) compared with those of control peptide in the presence of IL-15. Consistent with these results, the combination of full-length PAP with IL-15 also significantly increased IFN-γ responses in CD8+ T and NKT cells compared to IL-15 with or with control peptides (fig. S3). Overall, these results indicate that IL-15 significantly enhances tumor-specific CD8+ T and NKT cell responses of sip-T.

**Fig. 4.**
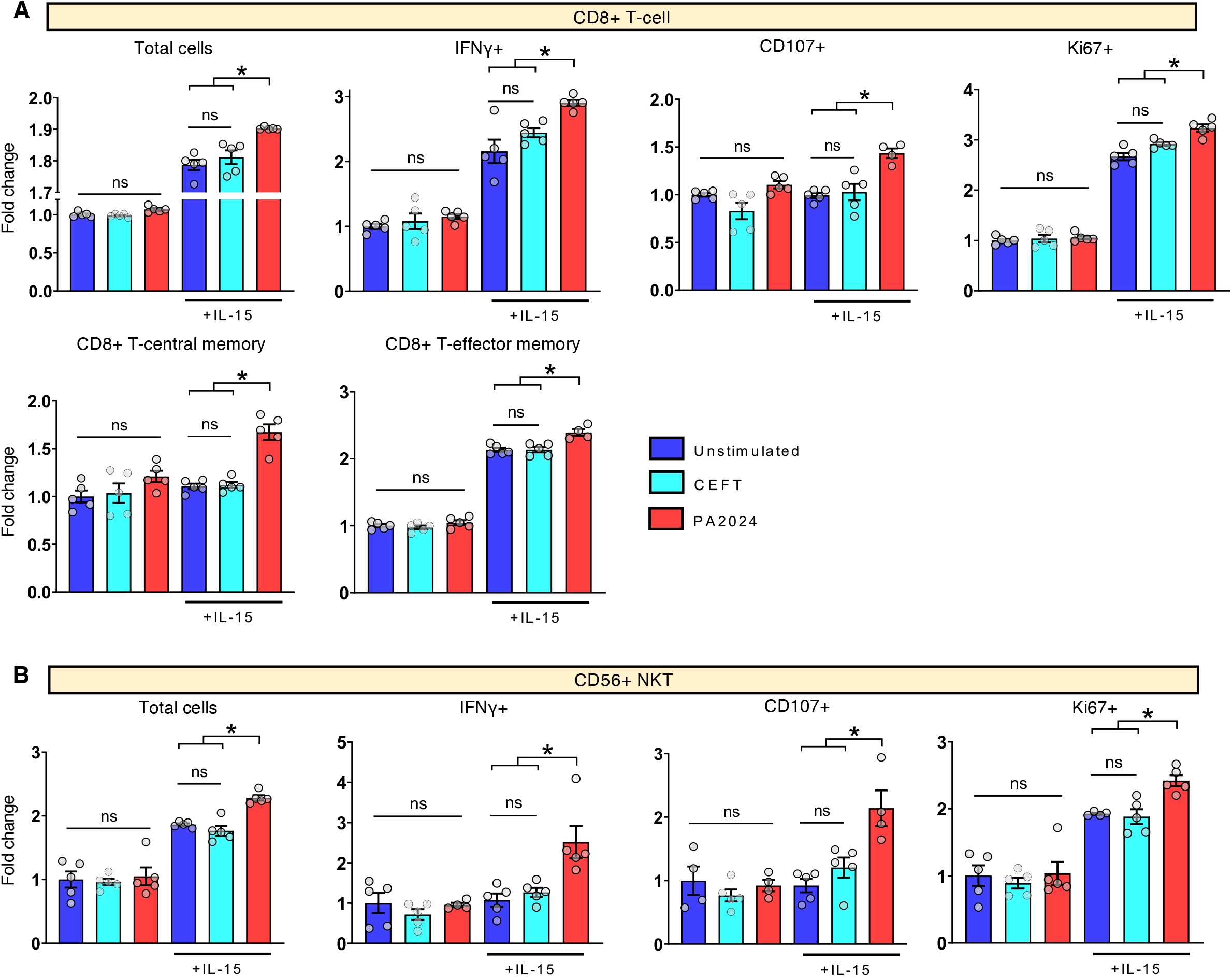
IL-15 expands tumor-specific effector T-cell subsets of sip-T. Sip-T from two individual mCRPC patients (n=5) were treated with PA2024 (25 µg/ml) or control peptide, CEFT (1 µg/ml) alone or in combination with IL-15 (50 ng/ml). Unstimulated controls received media only. IL-15 was added on day 0, 2, 4, 6 and cells harvested at day 7. Bar graphs (mean ± SEM shown) represent fold change relative to unstimulated controls. The data represents one of the two independent experiments **(A)** IL-15 treated sip-T showed significant expansion of IFN-γ+, CD107+, Ki67+, central memory (CCR7+CD45RA-), and effector memory (CD45RA-CCR7-) subsets of CD8+ T-cells following PA2024 but not CEFT treatments. **(B)** IL-15 treated sip-T exhibited a significant expansion of IFN-γ+, CD107+, and Ki67+ subsets of CD56+ NKT following PA2024 but not CEFT treatments. *, *P < 0.05* by 1-way ANOVA with Tukey’s post hoc test.

### CD8+ T cells and CD56+ NK/T cells contribute to IL-15-mediated increase of sip-T efficacy

IL-15 is a pleiotropic cytokine that can affect both innate and adaptive immune responses. IL-15 enhances the function, proliferation, and survival of NK cells and CD8+ T cells both of which play a key role in controlling tumors (*23*). Our initial studies with IL-15 showed significant proliferation and activation of both CD8+ T cells and CD56+ NK/T cells in sip-T (Fig. 2). In order to evaluate the contribution of each subset to the overall activation of sip-T, we performed depletion studies of both CD8+ and CD56+ subsets. The depletion resulted in successful removal of ≥90% and ≥93% CD8+ T-cell and CD56+ NK/T subsets respectively (data not shown). Supernatants from control or subset-depleted IL-15 treated sip-T were evaluated by ELISA (Fig. 5A). Depletion of either CD8+ T cells or CD56+ NK/T cells substantially significantly reduced the levels of IFN-γ, TNF-α, granzyme A, granzyme B, granulysin, sFAS, sFAS-L, and perforin compared to undepleted sip-T. Except for sFAS (which was significantly lower in CD8+ vs. CD56+ depletion), no significant differences were found in cytokine levels between CD8+ and CD56+ depleted fractions of sip-T, suggesting that both CD8+ T cell and CD56+ NK/T cells produce similar quantities of these cytokines in sip-T during IL-15 treatment. Depletion of both CD8+ T and CD56+ NK/T cell subsets further reduced cytokine levels, which were significantly lower compared with either of individual depletions (Fig. 5A).

**Fig. 5.**
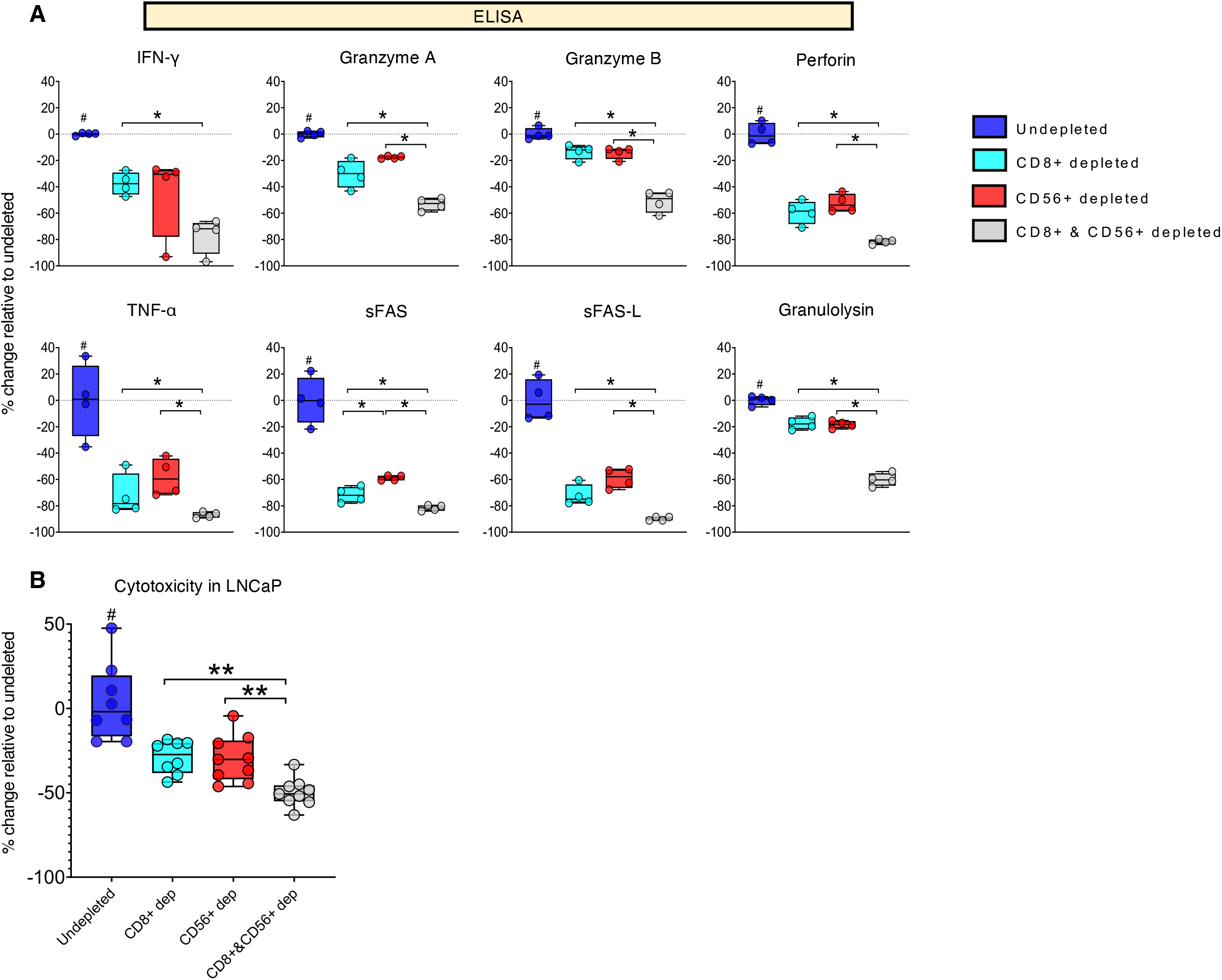
CD8+ T-cell and CD56+ NK/T cells are the primary source of IL-15-mediated enhancement of sip-T efficacy. CD8+ T-cell or CD56+ NK/T cell or both were depleted in sip-T from individual mCRPC patients. All depleted or undepleted (control) fractions were treated with IL-15 (50 ng/ml) on day 0, 2, 4 and 6 and harvested on day 7. **(A)** ELISA on cell supernatants (n=4 patients). The depletion of CD8+ T-cell or CD56+ NK/T cell or both significantly abrogated IL-15 mediated cytokine secretion (IFN-γ, granzyme A/B, perforin, TNF-α, sFAS, sFAS-L, and granulysin) from sip-T. **(B)** Cytotoxicity assay. IL-15 treated sip-T were washed and co-cultured for 24 hours with CFSE-labeled LNCaP (4:1). The depletion of either CD8+ T-cell or CD56+ NK/T cell or both significantly reduced IL-15 mediated cytotoxicity of sip-T compared with undepleted controls. #, *P < 0.*05 compared with any other groups; *, *P < 0.05*; **, *P < 0.01* by 1-way ANOVA with Tukey’s post hoc test.

To determine the role of each CD8+ T and CD56+ NK/T subset on the efficacy of sip-T, we performed cytotoxicity assays using control or depleted IL-15 treated sip-T. In line with our cytokine data (Fig. 5A), the depletion of either CD8+ T cell or CD56+ NK/T cell fractions significantly reduced IL-15 mediated enhancement in sip-T cytotoxicity in LNCaP tumors (Fig. 5B). Depletion of both CD8+ and CD56+ fractions further reduced IL-15-mediated sip-T cytotoxicity in LNCaP tumor cells. Taken together, these data suggest that CD8+ T cell and CD56+ NK/T cells are the key effector lymphocyte subsets that contribute to IL-15 mediated sip-T activation and augmentation of its direct anti-tumor efficacy.

### IL-15 unleashes sip-T from TGFβ mediated suppression of activation and function

Transforming growth factor beta (TGFβ) can promote prostate cancer growth and spread by causing immunosuppression, extracellular matrix degradation, epithelial to mesenchymal transition and angiogenesis, thereby promoting tumor cell invasion and metastasis (*24*). Studies have shown that TGFβ directly targets CD8+ T cells to inhibit the expression of key molecules involved in tumor cytotoxicity (*25*). Particularly relevant to mCRPC patients receiving sip-T, TGFβ is found at high levels in the prostate bone metastatic environment where it restrains optimal T cell development and function (*26*). To determine if IL-15 could rescue TGFβ-mediated suppression of sip-T activation and function, we performed in vitro assays with sip-T, treated with TGFβ alone or in combination with IL-15. Sip-T from four mCRPC patients was cultured with TGFβ (10 ng/ml) alone or in combination with IL-15 (50 ng/ml), followed by analysis with flow cytometry or a functional cytotoxicity assay (Fig. 6). TGFβ exposure did not significantly alter the percentage of CD8+ T and NK cells in sip-T, however we saw a significant reduction in CD56+ NKT cells compared with untreated control (data not shown). Importantly, the addition of IL-15 to TGFβ treated sip-T resulted in significant increases in all three lymphocyte subsets (Fig. 6A). TGFβ abrogated the activation (by CD107 expression) and proliferation (by Ki67 expression) of CD8+ T cell, CD56+ NKT, and NK populations of sip-T (Fig. 6, B-D), consistent with its known immunosuppressive effects (*27*). IL-15 significantly reduced TGFβ mediated suppression of sip-T, and partially or fully recovered proliferation (Ki67) in the three subsets; strikingly, the addition of IL-15 was able to induce a ∼2-to-4-fold increase in CD107 expression in the presence of TGFβ in both CD8+ T and NKT cell populations compared to untreated control (Fig. 6, B-D). We then sought to determine whether TGFβ had a functional impact on sip-T anti-tumor efficacy and if that could be countered by IL-15 treatment. Sip-T from four mCRPC patients was treated with TGFβ (5 or 10 ng/ml) alone or in combination with IL-15 (50 ng/ml), followed by co-culture with CFSE-labeled LNCaP and DU145 prostate tumor cells. TGFβ exposure resulted in significant suppression of sip-T mediated cytotoxicity in LNCaP and DU145 tumors (Fig. 6E). The addition of IL-15 overcame TGFβ-mediated suppression of sip-T cytotoxicity and significantly enhanced its anti-tumor efficacy against both tumor lines tested, at both concentrations of TGFβ (Fig. 6E). Overall, these data indicate that IL-15 can significantly reverse the immunosuppressive effects of TGFβ on sip-T, improving its activation/proliferation and anti-tumor cytotoxicity.

**Fig. 6.**
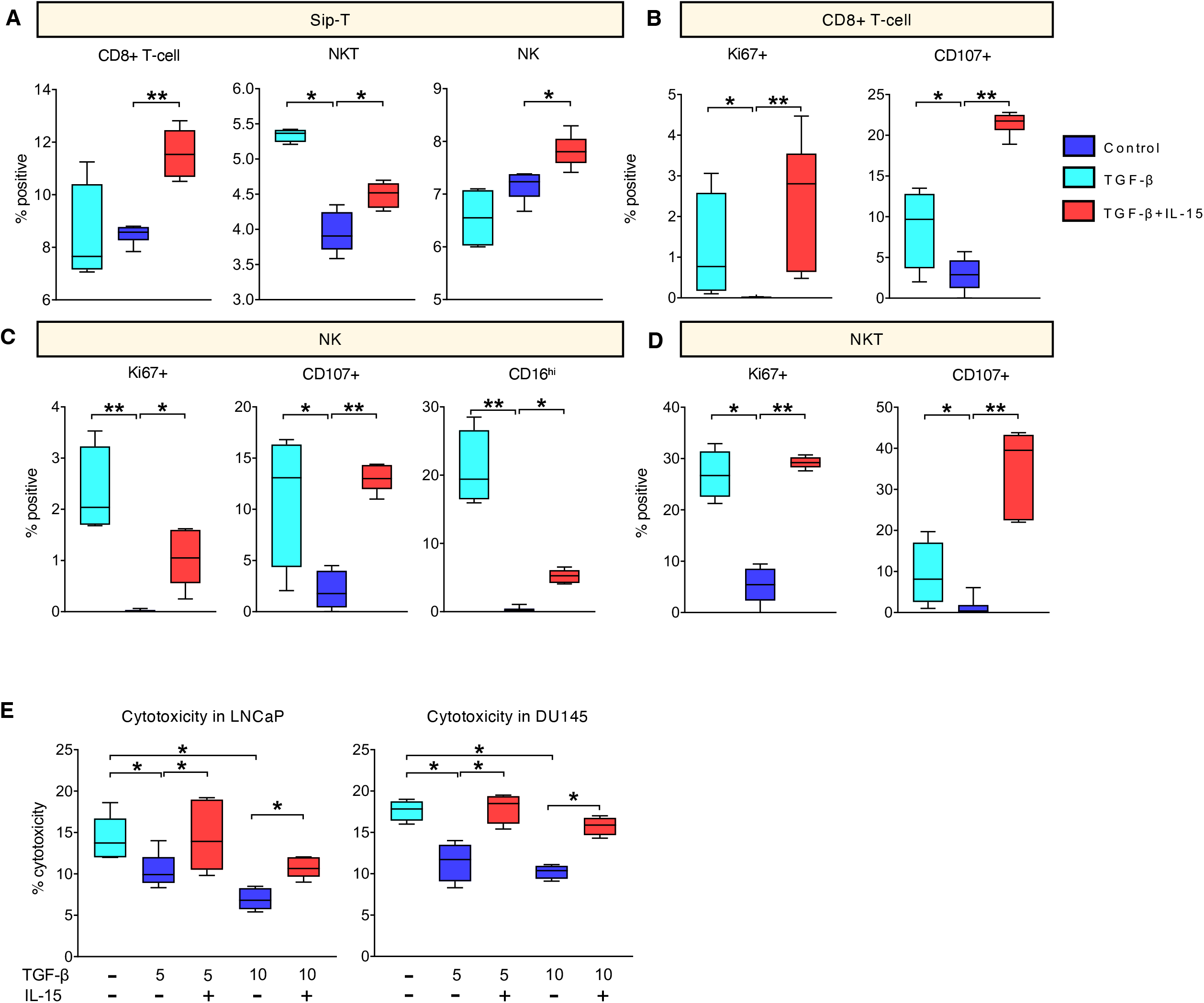
IL-15 overcomes TGFβ mediated suppression of sip-T activation and function. Sip-T from four mCRPC patients (n=4-6) were treated with TGFβ (5 or 10 ng/ml) alone or in combination with IL-15 (50 ng/ml) at day 0, 2, and 4 and harvested at day 5. **(A-D)** Assessment of proliferation (Ki67 expression) and activation (CD107 and CD16 expression) of effector sip-T (CD8+ T-cell, CD56+ NKT and NK cells) by flow cytometry. TGFβ (10 ng/ml) significantly reduced proliferation, activation, or both of effector sip-T; these effects were significantly rescued by the addition of IL-15. **(E)** Cytotoxicity assay. Cytokine treated sip-T were co-cultured with CFSE-labeled LNCaP or DU145 for 24 hours and analyzed by flow cytometry. TGFβ abrogated sip-T mediated cytotoxicity in LNCaP and DU145. IL-15 significantly rescued TGFβ mediated suppression of sip-T cytotoxicity. *, *P < 0.05*; **, *P < 0.01* by 1-way ANOVA with Tukey’s post hoc test.

### IL-15 significantly improves sip-T tumor growth inhibition

We next examined whether IL-15 treatment could enhance sip-T anti-tumor efficacy in vivo. NSG mice were engrafted with human prostate LNCaP tumors, and once established we adoptively transferred four doses of IL-15 treated or control (vehicle treated) sip-T with one-week intervals (Fig. 7A). To ensure uniformity of treatment, sip-T used in these experiments was generated from PBMCs from a single healthy donor utilizing PA2024 and following a protocol provided by Dendreon Pharmaceuticals LLC (see Methods). Two groups also received recombinant human IL-15 or vehicle subcutaneously at the time of sip-T administration; additional control groups received IL-15 or vehicle only, but no sip-T. We confirmed the presence of sip-T in mouse peripheral blood at day-7 after each sip-T administration (data not shown). As shown in Fig. 7B, treatment with sip-T but not IL-15 alone, significantly reduced tumor growth compared to untreated control. Concordant with our in vitro results, IL-15 treated sip-T significantly reduced tumor growth compared with sip-T only (over 2-fold) or the other control groups (over 3-fold; Fig. 7B).

**Fig. 7.**
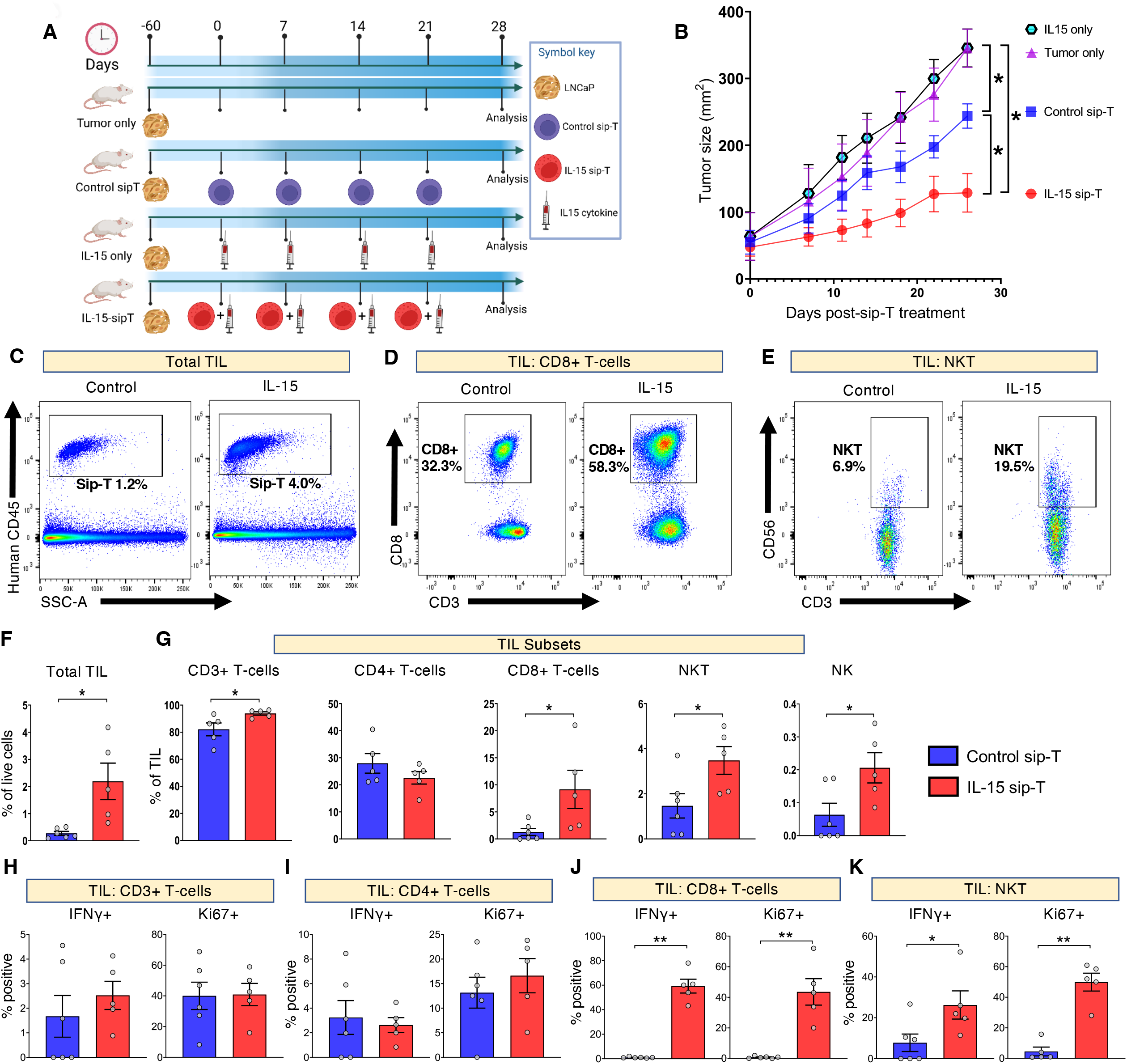
IL-15 significantly improves sip-T mediated suppression of tumor growth *in vivo*. Male NOD/SCID/IL2R gamma (null; NSG) mice were implanted subcutaneously with 3×10^6^ LNCaP cells. Once the average tumor size reached 50-100 mm^2^, mice were divided into 4 groups, i.e., tumor only, control sip-T, IL-15 only, and IL-15 sip-T. Sip-T groups received four injections of either control or IL-15 treated sip-T (5×10^6^ cells intraperitoneally), seven days apart. IL-15 sip-T and IL-15 only groups also received four injections of rhIL-15 (1 µg subcutaneously), seven days apart. Mice were euthanized and tumors analyzed, *ex-vivo*, on day 28 post-initiation of sip-T treatment. **(A)** Schematic illustration of the mouse experiment layout **(B)** Graphs are from a representative experiment of two performed. Tumor size is represented as mean ± SEM, with cohorts of 4-6 mice per group. **(C-E)** Analysis of tumor-infiltrated lymphocytes (TILs) by flow cytometry. Representative dot plots of major immune cell subsets affected by IL-15 treatment. **(F-K)** Bar graphs of sip-T analysis in TILs (mean ± SEM). IL-15 treated sip-T showed significantly higher expansion, activation (IFN-γ expression) and proliferation (Ki67 expression) of CD8+ T-cell and CD56+ NKT and expansion of CD56+ NK cells compared with respective controls. *, *P < 0.05* by Mann-Whitney.

Tumors were resected and analyzed on day 28 post-administration of the first sip-T injection. Tumor-infiltrated lymphocyte (TIL) analysis by flow cytometry revealed that the frequency of human CD45+ cells (i.e., sip-T) was 2 to 15-fold higher (greater than 8 on average) in tumors from mice that received IL-15 treated sip-T (IL-15 sip-T) compared to control sip-T (Fig. 7 C, F). Infiltrating total T cells (CD3+), CD8+ T cells, CD56+ NKT and NK cells - but not CD4+ T cells - were significantly higher in tumors from mice that received IL-15 sip-T compared with those of control sip-T (Fig. 6, D-E, G). Treatment with IL-15 sip-T significantly increased activation (by IFN-γ+) and proliferation (by Ki67+) of tumor-infiltrating CD8+ T and NKT cells, but not total T cells (CD3+) or CD4+ T cells compared to control sip-T (Fig. 7 H, I).

Analysis of peripheral blood on day 28 post-administration of the first sip-T injection revealed no significant differences in circulating sip-T (total CD45+ cells) frequencies between the two groups; further subset analyses showed a significant increase in peripheral CD8+ T cell with a concomitant decrease in CD4+ T cells and no significant change in NKT and NK cell subsets (fig. S4). Strikingly, however, IL-15 treatment significantly increased IFN-γ+ and Ki67+ CD8+ T cell and CD56+ NKT cell populations, while no significant changes were seen in total CD3+ T and CD4+ T cells compared with control (fig. S2 B-F).

### IL-15 reshapes the sip-T and tumor microenvironment transcriptome to enhance anti-tumor immunity

To further characterize the effects of IL-15 treatment on sip-T mediated tumor immunity, we investigated IL-15-dependent gene expression changes by performing RNA-seq on tumors resected from NSG mice on day 28 post-initiation of sip-T treatment. RNA-seq data showed a differential upregulation of a larger set of genes (470 genes in total) in tumors from IL-15 sip-T group compared with only 243 genes in the control sip-T or 256 genes in tumor only groups (Fig. 8A). As treatment with IL-15 alone had no effect on in vivo tumor growth (Fig. 7B) and as LNCaP tumors do not express IL15Rβ or the common γ chain, with little to no expression of IL15Rα (*28*) we did not include these tumors in our analyses. The functional characterization of the differentially expressed genes between the IL-15 treated and control sip-T groups was performed using Gene Ontology (GO) term enrichment analysis as described previously (*29, 30*). The expression of 13336 genes was equally regulated between IL-15 treated and control sip-T groups (Fig. 8A). Pathway analysis of IL-15-targeted upregulated genes showed enrichment of signature pathways involved in lymphocyte differentiation, activation, proliferation, cytotoxicity, lysosome/autophagosome, and antigen binding, processing, and presentation (Fig 8B). Of 470 genes differentially upregulated by IL-15 treatment, Fig. 8C shows 28 genes relevant to sip-T anti-tumor immunity that were found in common across the enriched pathways in Fig. 8B. Detailed information for significantly upregulated genes and enriched pathways is provided in table S4. These genes encompass signature molecules for effector lymphocyte development, differentiation, proliferation, and activation (*CD3d*, *CD8a*, *CD8b*, *IL2RA*, *IL2RB*, *IRF5*, *JAK3*, *NKG2*, *CD40L*, *SLAMF1*), lymphocyte trafficking (*CCL3*, *CCL5*, *IL18*), antigen recognition, binding, processing, and presentation (*HLA-B*, *HLA-DR*, *HLA-DQR1*, *ULBP1*, *NLRC3*, *MOAP1*) and anti-tumor /cytotoxic/phagocytic activity (*FASLG*, *GZMA*, *GZMB*, *GZMH*, *GNLY*, *IFNG*, *TNFSF10*, *TNFS12*, *UNC13D*). We further confirmed the expression of several key genes found upregulated in RNA-seq data (with IL-15 treatment), along with additional TME-relevant genes by qPCR (Fig. 8D). We found significant upregulation of genes (*TNFA*, *MKI67*, *IFNG*, *CD107*, *HLA-A/B/C*, *IRF5*, *CCL5*, *GZMB*, *FASL*, *NKG2*) known to play a role in lymphocyte proliferation and activation, antigen recognition and presentation, and anti-tumor/cytotoxic activities (*29, 30*) in the IL-15 treated group. Importantly, we found that genes known to promote tumor growth and/or immune-suppression (*RANKL*, *B7H3*, *PDL1*/*CD274*, *CCL2*, *CXCL8*, *CXCL9*, and *TGFB*) (*24, 31–33*) that were significantly *upregulated* in control sip-T treated tumors were significantly *downregulated* in tumors treated with IL-15 sip-T (Fig. 8D). Overall, these results show that IL-15 favorably impacts the sip-T and TME transcriptome by significantly upregulating key pathways involved in lymphocyte effector activity, and reverses sip-T-induced expression of immunosuppressive genes such as TGFβ.

**Fig. 8.**
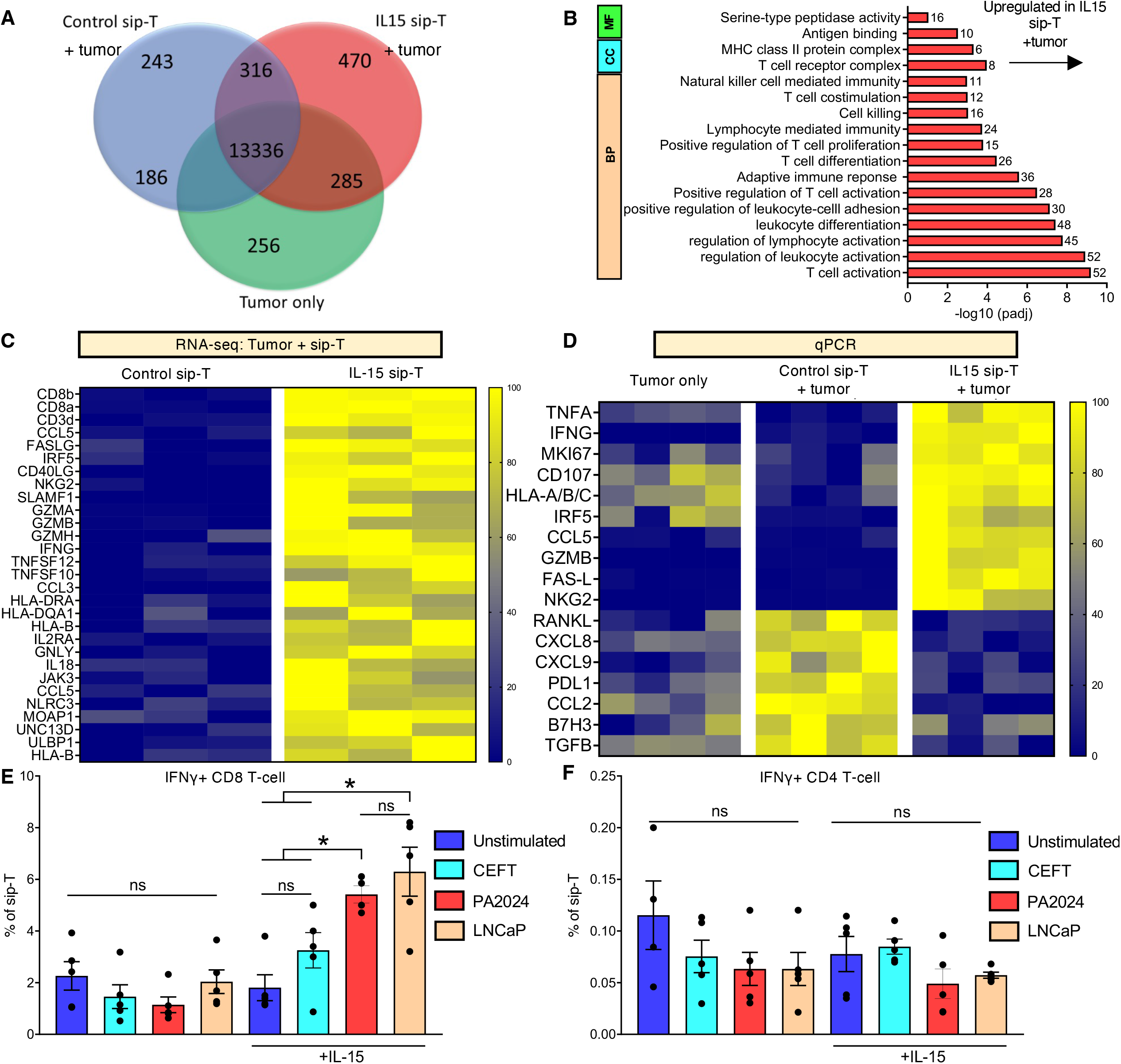
*Ex-vivo* functional analysis of IL-15 treated sip-T. Tumors resected from NSG mice (described in Figure 7) were analyzed by RNA-seq and qPCR. (A-C) Bulk RNA-seq of sip-T. **(A)** Venn diagram of differentially expressed genes among IL-15 sip-T, control sip-T and tumor only groups. **(B)** Differential gene pathway analysis of RNA-seq between tumors from IL-15 and control sip-T groups. IL-15 sip-T showed a significant enrichment in pathways involved in lymphocyte activation, proliferation, lymphocyte-mediated cytotoxicity, downstream antigen receptor signaling, lysosome/autophagosome, secretory/zymogen granules, and antigen and cytokine binding. **(C)** Heatmap displaying differential gene expression of activation/effector markers (by RNA-seq) showed that many of these genes were upregulated in mouse tumors which received IL-15 treated sip-T **(D)** Heatmap displaying differential gene expression of activation/effector markers (by qPCR) further confirmed that many of these genes were upregulated in IL-15 sip-T tumors. Some significantly downregulated genes following IL-15 treatment are also shown in the lower panels. **(E)** splenocytes (n=4-5) from NSG mice (described in Figure 7) were stimulated with LNCaP lysates (50 µg/ml), PA2024 (25 µg/ml) or control peptide, CEFT (1 µg/ml) for 24 h. Unstimulated controls received media only. Bar graphs (mean ± SEM shown) represent percent positive of total sip-T. Splenocytes from IL-15 sip-T but not control sip-T mice showed significant expansion of IFN-γ+ CD8 T-cells following LNCaP or PA2024 stimulations as compared with CEFT or untreated controls. *, *P < 0.05* by 1-way ANOVA with Tukey’s post hoc test.

### IL-15 expands tumor specific cytotoxic T cell subsets of sip-T in vivo

Finally, we sought to determine if treatment with IL-15 could enhance responses of tumor-antigen specific lymphocyte effector populations. To test this, we prepared splenocytes (on day 28 post-initiation of sip-T treatment) from NSG mice which received control or IL-15 treated sip-T. Splenocytes were treated with prostate tumor-specific antigens (i.e., PA2024 or LNCaP lysate), as well as control peptide (CEFT) or left untreated (i.e., media alone) for 48 h. In accordance with our in vitro data (Fig. 4 and S3), splenocytes from IL-15 sip-T but not control sip-T mice showed significant expansion of IFN-γ+ CD8 T-cells following LNCaP or PA2024 stimulation as compared with CEFT or untreated controls (Fig. 8E). Exposure to any of these antigens did not significantly alter CD4+ T-cell activation (Fig. 8F), suggesting that IL-15 treatment may preferentially help priming of antigen-specific CD8+T-cells in vivo. Our data suggest that IL-15 not only non-specifically expands and activates CD8+ T-cells (Fig. 7D, G, J), but acts to augment tumor antigen-specific responses in these cells ex-vivo. Overall, these results indicate that IL-15 significantly enhances tumor-specific CD8+ T cell responses of sip-T.

## DISCUSSION

Sipuleucel-T (sip-T) is the first and only therapeutic cellular therapy approved by the FDA for use in men with metastatic castration-resistant prostate cancer (mCRPC) (*3, 4*). Here, we present the first study characterizing the phenotype of sip-T in detail using high dimensional mass cytometry (CyTOF), as well as the first data showing the efficacy of sip-T in preclinical models of prostate cancer and improvement with cytokine treatment. Despite sip-T often being described as a dendritic cell (DC) or antigen presenting cell (APC) therapy, here we show that T cells (both CD4+ and CD8+) typically constitute the vast majority of sip-T, followed by B-cells, NK cells, NKT cells and monocytes in addition to a smaller proportion of dendritic cells, ILCs, and MDSCs. We observed considerable variability in the percentage of B cell and monocyte/macrophage populations in the sip-T product, albeit in the context of a relatively small sample size. Interestingly, patients treated with sip-T have been found to have elevated levels of IgG against multiple secondary tumor antigens (e.g., PSA, hK2/KLK2, KRAS, etc.) compared to controls, and significantly improved survival associated with those patients with IgG responses to PSA and galectin-3/LGALS3 (*34*). Our study did not examine humoral responses to sip-T, as we focused on characterizing the sip-T product itself and these humoral responses in patients have been well characterized in the past (*35*).

Based off promising results from our previous clinical trial of sip-T in combination with IL-7 (*2*), we characterized the effects of other clinically promising major γC cytokines (*36*) on sip-T effector function. We showed that treatment of sip-T with IL-15 significantly improved the efficacy of sip-T compared with IL-7 and IL-21, both of which have been tested clinically in recombinant and other forms (*12, 37*). These cytokines belong to the four α-helix bundle family which are endogenous homeostatic molecules that can stimulate the proliferation and activation of T cells, and augment T cell migration from blood into lymph nodes and peripheral organs (*9, 10*). Treatment with IL-15 significantly expanded effector lymphocyte populations compared to control and other cytokines, and we saw significant increases in CD107 expression as a marker of immune cell degranulation, as well as prostate tumor cell cytotoxicity. IL-15 treatment favorably altered the transcriptional profile of sip-T, increasing lymphocyte activation, antigen presentation, migration, and tumor cytotoxicity. IL-15 has the potential to broadly stimulate lymphocyte subsets, however we showed that IL-15 augmented antigen-specific activation of sip-T.

With respect to major effector populations within sip-T, our depletion studies support the key roles of CD8+ and CD56+ lymphocyte subsets in tumor cell cytotoxicity and production of mediators of tumor lysis/suppression. This, however, does not discount the potential role of APCs contained within sip-T that may have additional beneficial effects in vivo. As the bone TME in prostate cancer has relatively high TGFβ expression (*26*), we exposed sip-T to clinically relevant levels of TGFβ with or without IL-15 treatment and found that IL-15 can reverse the immunosuppressive effects of TGFβ on sip-T, in terms of proliferation, degranulation, and tumor cytotoxicity. Importantly, these data suggest the addition of IL-15 to sip-T may be a promising strategy to improve efficacy in prostate cancer patients with bone dominant disease.

Our mouse prostate tumor studies of human sip-T treatment - the first to our knowledge - showed that treatment of sip-T with IL-15 significantly improved anti-tumor efficacy against human prostate tumor xenografts. IL-15 treatment resulted in significantly more tumor-infiltrating leukocytes, with an approximately 4-fold increase in CD8+ T cells and a striking 59-fold increase in the percentage of IFNγ+ CD8 T cells. CD8+ T cell, NKT or NK cell infiltration and activation within the tumor are strongly correlated with improved clinical prognosis (*11, 13, 38*). In accordance with previous reports in other experimental settings (*11, 39, 40*), we found a significant expansion and activation of CD8+ T cell and CD56+ NKT/NK subsets of sip-T in the tumor microenvironment following IL-15 treatment. In fact, IL-15 is known to protect NKT cells from inhibition by tumor-associated macrophages and enhance anti-metastatic activity (*40*). RNAseq data from these in vivo studies revealed IL-15 mediated reversal of immune suppression induced by treatment with sip-T, which is in line with a previous study showing IL15 reversal of TGFβ mediated immune suppression (*41*). Given that low levels of IL-15 have been associated with poor prognosis in cancer patients (*13*), the addition of IL-15 to sip-T may be a promising approach to improving clinical outcomes in mCRPC patients.

To date, sip-T remains the only FDA-approved immunotherapy for genomically unselected (i.e. MSI-S, <10 mut/MB) mCRPC patients, and strategies to improve its clinical efficacy are warranted. Human recombinant IL-15 has been used in recent clinical trials, in combination with other immunotherapies, including emtuzumab (anti-CD52) (NCT02689453), ipilimumab and nivolumab (NCT03388632), mogamulizumab (anti-CCR4) (NCT04185220), and obinutuzumab (anti-CD20) (NCT03759184) (*42*). Our CyTOF data indicates that various sip-T subsets (e.g., CD4+ and CD8+ T cell and monocytes) exhibit high expression of immune checkpoint molecules including PD-1, PD-L1, ICOS and TIM3. Thus, IL-15 treatment of sip-T in combination with ICB therapy could offer an attractive therapeutic implication in mCRPC patients. We also propose that the insufficient expansion of CD8+ T cells by ICB therapy alone may potentially be overcome by combining IL-15 treatment with sip-T therapy in mCRPC patients. However, some limiting factors for IL-15 application in the clinic include its short half-life and low efficacy in vivo. IL-15 superagonists present an advantage over rIL-15 in terms of their longer half-life and immune modulatory effects, including the expansion of NK and CD8+ T cells, without inducing significant toxicity (*42*). The four main soluble and structurally distinct IL-15 superagonists include heterodimeric IL-15 (*43*), receptor-linker-IL-15 (RLI) (*44*), IL-15/IL-15Rα-Fc (*45*), and N-803 (formerly ALT-803) (*37*). Whether IL-15 superagonists vs. rIL-15 translates to better sip-T efficacy remains to be determined and requires further study.

Limitations of our study include our small sample size (n=14 from 11 individual patients), and subsequently we found high inter-individual variability in the composition of sip-T subsets, which precluded any correlations between the immune subsets in sip-T and clinical outcomes, and indeed the study was not powered to determine such correlations. Also, it was not possible to exclude the potential effects of a freeze-thaw cycle on sip-T composition given the logistical challenges of sample collection over time and weighed against the benefits of batched analyses, although these tend to be more pronounced in the myeloid component (*46*).

Overall, this study is the first to fully characterize the sip-T cellular composition by CyTOF, to evaluate the effects of cytokines on sip-T, and then to utilize *in vivo* modeling of sip-T with engrafted human prostate tumors. Our data indicate that IL-15 greatly enhances anti-tumor efficacy and immunity of sip-T against prostate tumors, with CD8+ T cell and CD56+ NKT/NK cells identified as key effector populations. Transcriptomic analyses revealed IL-15 mediated reversal of sip-T induced immunosuppression within the TME, which has not previously been described. Our studies thus support the clinical investigation sip-T in combination with rIL-15 or IL-15 analogs as a promising strategy to augment anti-tumor immunity and potentially improve clinical outcomes in patients with mCRPC.

## MATERIALS AND METHODS

### Collection of sipuleucel-T

Sip-T samples from mCRPC patients were collected after their verbal and written consent under the IRB approved banking protocol, IRB#201411135. All demographic and clinical characteristics of these patients were also documented. Briefly, the infusion bag and tubing were disconnected from the patient after sip-T administration and carried to the lab on ice. Both the infusion bag and tubing were washed with ice cold 50 ml phosphate buffered saline (PBS) and sip-T were pelleted by centrifugation for 5 min at 300 G. The cells were washed twice with PBS and frozen in 90% fetal bovine serum (Peak Serum) with 10% DMSO in Mr. Frosty freezing container at -80°C for 24 hours and then transferred to liquid nitrogen until use.

### CyTOF analysis

Sipuleucel-T from eleven mCRPC patients (n=14 samples) were analyzed using high dimensional single cell analysis by CyTOF. Cryopreserved sip-T were thawed in RPMI complete (RPMI 1640 (Gibco Life Technologies) +5% human AB serum (Sigma) +2 mM L-glutamine +100 units penicillin-streptomycin +25 mM HEPES buffer +1x non-essential amino acids (NEAA). Cells were washed and resuspended at 2×10^6^ cells/mL, and rested for 60 min at 37°C, 5% CO_2_ before use in mass cytometry (CyTOF).

A complete list of all antibodies used for CyTOF is provided in Table S1. Briefly, sip-T were resuspended in PBS as 1-3×10^6^ cells per sample for staining. Cells were stained in 5 μM cisplatin for 3 min at room temperature, followed by washing with Cy-FACS buffer (PBS, 0.1% BSA, 0.02% NaN3, 2 mM EDTA) twice. Cells were then blocked with FcR blocking reagent (Biolegend) on ice for 10 min followed by addition of surface-antibody cocktail in the mixture for another 60 min. After incubation, cells were washed twice with PBS and then fixed/permeabilized with eBioscience Fixation/Permeabilization reagent (Invitrogen) for 30 min on ice and stained for 60 min with the intracellular antibody cocktail. Cells were then fixed with 4% PFA for 20 min on ice, washed twice with PBS and then stained with DNA intercalator (200 μl per 10^6^ cells). Cells were then run, and data acquired on a Fluidigm CyTOF 2 Mass Cytometer. Data were exported to FlowJo (V10.8.1) or OMIQ for further analysis. All analyses were then made on live, single CD45+ cells followed by manual gating based on lineage-marker expression to characterize immune populations. To spatially visualize sip-T subsets, a UMAP spatial heatmap algorithm analysis was applied to CD45 + cells using OMIQ.

### Antigen-specific T cell response detection in sip-T

Antigen-specific T cell responses of sip-T were detected as described previously (*2*). Briefly, fresh sip-T (n=5 samples from two individual patients) were resuspended in RPMI complete and plated at 2×10^6^ cells/mL in 24-well ultra-low attachment plate (Corning Life Sciences). Sip-T was stimulated with 25 μg/ml PA2024 (Dendreon) or 1 ug/mL CEFT (Cytomegalo virus, Epstein-Barr virus, Influenza virus, and Tetanus toxin peptides; negative control). Cells were then stimulated with 50 ng/ml rhIL-15 (BioLegend) on day 0, 2, 4, 6. The cells were harvested for flow cytometry analysis on day 7. The assay was repeated at least twice.

### Cytokine stimulation assays

Fresh or freshly thawed (cryopreserved cells thawed in RPMI complete) sip-T was resuspended in RPMI complete and plated at 1-2×10^6^ cells/mL in 24-well ultra-low attachment plate (Corning Life Sciences). All in vitro assays were performed using sip-T from at least 3 individual mCRPC patients (to account for any inter-individual variability) and each sample was split into unstimulated and stimulated fractions. For cytokine stimulations, each sample was treated with recombinant human IL-7, IL-15, or IL-21 (50 ng/ml) for 5-7 days. Cytokines were replenished on alternative days (i.e., day 0, 2, 4, 6). Each in vitro stimulation assay was repeated at least twice. For some assays with freshly thawed sip-T, a low dose rhIL-2 (<20 IU/ml) was added to all wells including control for better revival though all such assays were repeated in the absence of IL-2.

### Flow-based cytotoxicity assays

Sip-T left untreated (control sip-T) or treated with IL-15 or other cytokines (see cytokine stimulation assays) were used for cytotoxicity assays. Trypsinized tumor cells were counted and stained with carboxyfluorescein diacetate succinimidyl ester (CFSE, 1 μL/mL; BioLegend) following the manufacturer’s protocol and cultured overnight. Cytokine-treated sip-T were washed twice and added to CFSE-labeled target tumor cells at indicated effector to target (E:T) ratios (typically 4:1) and cultured overnight. Samples were stained with 7-AAD (5 μL/1 × 10^6^, BioLegend) to identify dead cells. Percent cytotoxicity was the fraction of cells that stained positive for both CFSE and 7-AAD.

### Incucyte-based cytotoxicity assays

IL-15 treated, or control sip-T were prepared as described in ‘cytokine stimulation assays’ before starting this assay. Nuclight Red (Sartorius)-labeled LNCaP or DU145 were seeded at 12,000 cells/well in 100 µl phenol red-free RPMI in a 96-well plate and incubated for 18 hours. IL-15 treated or control sip-T (see cytokine stimulation assays for details) were washed twice, resuspended in phenol red-free RPMI containing Cytotox Green (1/4000 diluted, Sartorius). Tumor cells were washed twice with phenol red-free RPMI. Sip-T was then added to the Nuclight Red-labeled target tumor cells at indicated effector to target (E:T) ratios (typically 4:1). Cell death was measured via the Cytotox Green probe, and live-cell analysis was measured using Nuclight Red. Cell death was analyzed using IncuCyte FLR imaging system software (Sartorius). For quantification of death, Cytotox Green florescence was normalized to total counts for each respective well. To calculate the relative cytotoxicity (fold change) at each timepoint, the values of cell death for IL-15 treated sip-T were normalized to those of control sip-T sip.

### Flow cytometry

All surface and intracellular antibodies used for flow cytometry are enlisted in Table S3. Sip-T or single cell suspensions of tumor-infiltrating lymphocytes (TILs) were resuspended in PBS as 1-3×10^6^ cells/ml and stained with fixable Zombie Aqua viability dye (BioLegend) as per manufacturer’s recommendation. Cells were washed twice with FACS buffer (PBS containing 0.5% BSA and 3 mM EDTA), blocked with FcR blocking reagent (BioLegend) on ice for 10 min and then incubated with surface antibody cocktail for 30 min in dark. Intracellular staining was performed after surface staining where applicable. For intracellular staining, the cells were fixed/permeabilized using the eBioscience Foxp3 Staining Buffer Set (Invitrogen) following manufacturer’s protocol and then stained with intracellular antibody cocktail for 30 min on ice. FCS Data were acquired on BD Fortessa X-20 (BD Biosciences) within 1-2 days following antibody staining and analyzed using FlowJo software (v10.8.1).

### In vivo studies

Approximately, 9-10 weeks old male NOD/SCID/IL2R gamma (null) (NSG; No. 005557, *NOD.Cg-Prkdc^scid^ Il2rg^tm1Wjl^/SzJ*) mice from the Jackson Laboratory were used in these studies. Mice were maintained in the Washington University facilities (St. Louis, MO) following the guidelines of the Division of Comparative Medicine. To evaluate the effect of IL-15 treated sip-T on in vivo tumor growth, LNCaP tumor cells (3 × 10^6^) were inoculated subcutaneously in mice in 200 µl RPMI with 50% matrigel (BD Biosciences). Once average tumor sizes reached 50-100 mm^2^, mice received four injections of IL-15 treated, or control sip-T (5×10^6^/100 µl in PBS, intraperitoneally) with one-week intervals. rhIL-15 (1 µg) was also injected subcutaneously at the time of sip-T injection. All control mice were sham dosed with vehicle (PBS). On day 7 post-last-sip-T injection, mice were humanely euthanized by carbon dioxide inhalation followed by cervical dislocation and tissues collected for analysis.

Single cell preparations of tumor-infiltrating lymphocytes were made by mechanically disrupting and enzymatically digesting tumors using gentleMACS™ (Miltenyi Biotec). The digestion media contained collagenase IV and dispase II (1 mg/ml each; Thermo Fisher) in DMEM media. Due to the requirement of a large number of sip-T for animal studies, HLA-matching for each injection into the same animal, and limited availability of patient sip-T, we prepared in house sip-T for adoptive transfer experiments following the Dendreon guideline. Briefly, PBMCs from human volunteers isolated by lymphocyte separation medium (Corning) were resuspended in AIM-V (Gibco) as 10^7^ cells/mL in ultra-low attachment flasks (Corning). Cells were incubated with 10 mg/mL PA2024 (kindly provided by Dendreon) for 36-42 hours. Cells were then treated with IL-15 (50 ng/ml) or vehicle (PBS) for 24 hours before injecting into mice.

### RNA sequencing (RNA-seq) analysis

Sample RNA was isolated using TRizol (Invitrogen) following manufacturer’s recommendation. Illumina sequencing libraries were constructed by Novogene Co., Ltd., which also performed pair-end sequencing with Illumina HiSeq3000. Differential expression analysis of two groups was performed using the DESeq2 R package (1.20.0). The resulting P-values were adjusted using Benjamini and Hochberg’s approach for controlling the false discovery rate. Genes with an adjusted P-value ≤0.05 found by DESeq2 were assigned as differentially expressed. Gene Ontology (GO) enrichment analysis of differentially expressed genes was implemented by the cluster Profiler R package, in which gene length bias was corrected. GO terms with corrected P-value less than 0.05 were considered significantly enriched by differential expressed genes.

### Real-time RT-PCR analysis

Sample RNA was isolated using TRizol (Invitrogen) following manufacturer’s recommendation. RNA concentrations were determined using NanoDrop 2000 (Thermo Fisher Scientific). High-Capacity cDNA Reverse Transcription Kit (Thermo) was used to convert RNA to cDNA via the manufacturer’s protocol. cDNA was amplified with Power SYBR Green Master Mixi (Thermo) via the manufacturer’s protocol using human primer-sets enlisted in Table S5. A StepOnePlus™ Real-Time PCR System (Thermo) was used to quantify mRNA expression via the 2^-ΔΔCt^ analysis method as described previously (*47*). Gene expression was normalized to GAPDH for each sample individually and then normalized to mean of control group across the dataset.

### Estimation of cytokines and secretory molecules

Cytokines secreted in the culture supernatant of cytokine-stimulated sip-T were measured by a LEGENDplex Human CD8/NK Panel kit (BioLegend) following manufacturer’s protocol. The data of the LEGENDplex multi-analyte flow assay were collected on a BD Fortessa X-20 (BD Biosciences). ELISA for IFN-γ and IL-10 were also performed with human ELISA MAX Deluxe Sets (BioLegend) to verify the multiplex ELISA results. Cytokine concentrations in each sample were calculated against a standard curve with known concentrations of recombinant cytokine.

### Statistical Analyses

All statistical analyses were performed using GraphPad Prism software (v9). All data are representative of at least two independent experiments (n=3 or more), unless specifically noted. Means and SEM were calculated. Paired Student t tests or nonparametric Mann-Whitney U tests were used for comparison between two groups in each experiment. One-way ANOVA (including the Turkey’s post hoc) was used to compare more than two groups. A p-value of less than 0.05 was considered statistically significant.

### Study approval

Sip-T samples from mCRPC patients were collected after their verbal and written consent under an IRB approved banking protocol (#201411135). All animal studies were performed in accordance with approved Washington University (St. Louis, MO) and NIH Institutional Animal Care and Use Committee guidelines under an approved protocol (No. 20-0383).

## List of Supplementary materials

Fig. S1 to S4

Tables S1 to S5 (Excel files)

## Acknowledgments

We thank the Alvin J. Siteman Cancer Center at Washington University School of Medicine and Barnes-Jewish Hospital in St. Louis, MO., for the use of the Bursky Center for Human Immunology and Immunotherapy Programs Immunomonitoring Laboratory which provided CyTOF services. The Siteman Cancer Center is supported in part by an NCI Cancer Center Support Grant #P30 CA091842.

## Author contributions

RKP conceived the idea for this study. MAS and RKP designed all the experiments. MAS, BP, and KK performed all experiments and collected the data. MAS and RKP analyzed all the data. AB, BP, and KK helped with the collection and processing of patient samples. BVT helped with the Incucyte assays. MB, DT, SN, VT and TF provided support and materials for in vitro and in vivo assays. MAS and RKP drafted the manuscript. All authors reviewed and contributed to the manuscript.

---

**Fig. S1.**
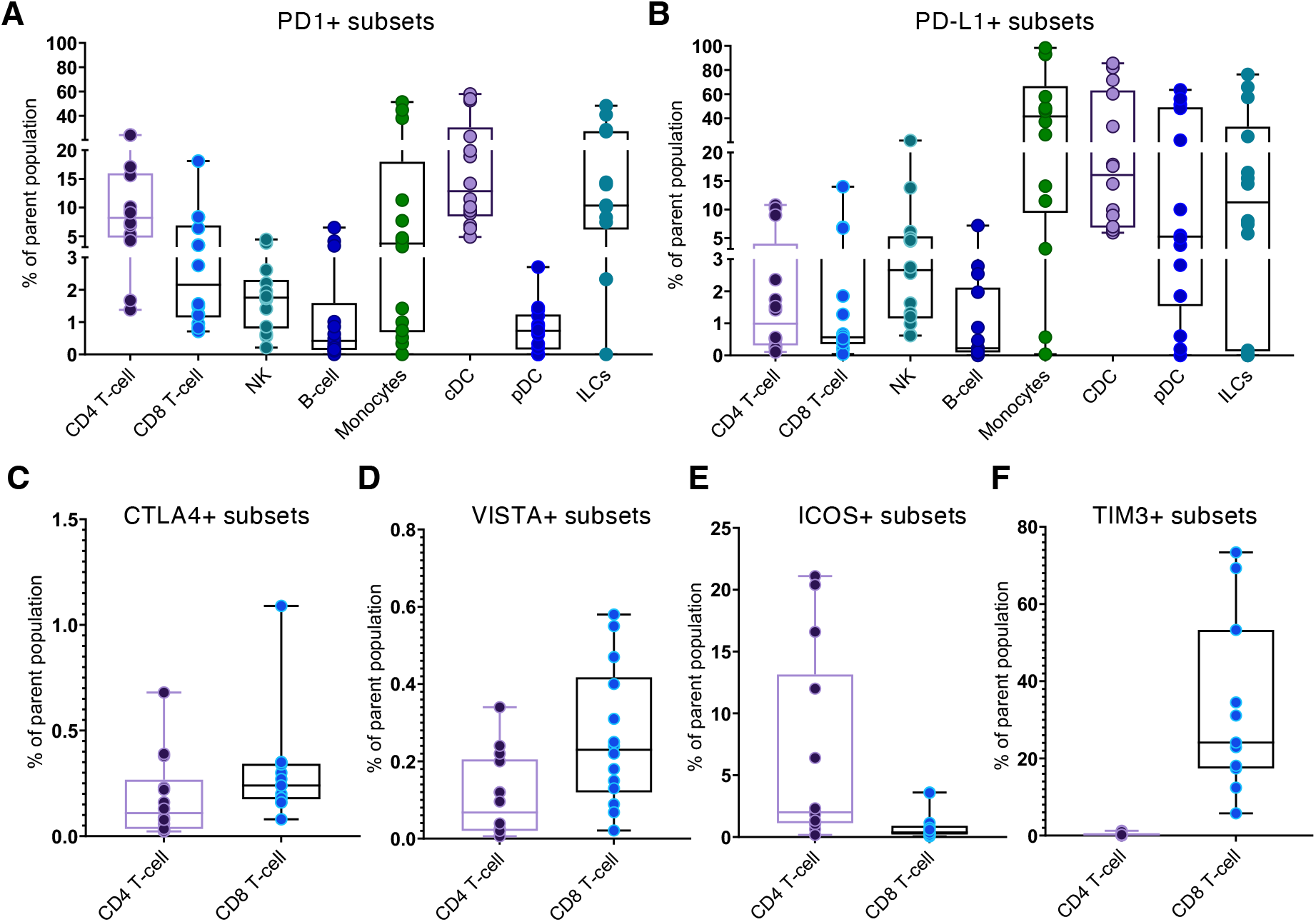
CyTOF analysis of various immune checkpoint molecules in sip-T subsets. Fourteen sip-T samples from 11 individual mCRPC patients were analyzed by mass cytometry (CyTOF). **(A-F)** the expression of various immune checkpoint molecules presented as % positive of the parent population.

**Fig. S2.**
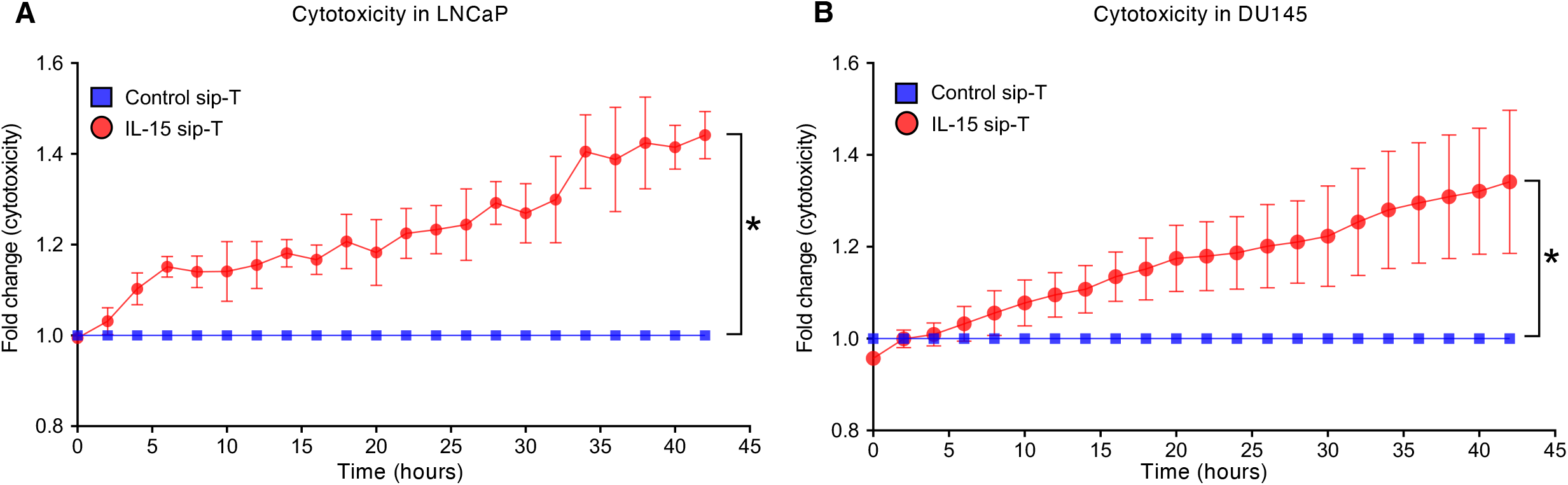
IL-15 treatment of sip-T significantly improves their cytotoxicity (determined by incucyte) against prostate tumors. Sip-T from individual mCRPC patients were treated with IL-15 (50 ng/ml) or control (media only) on day 0, 2, 4, 6 and then washed thoroughly and used for cytotoxicity assay at day 7. Incucyte-based cytotoxicity assay. IL-15 treated sip-T co-cultured with Nuclight Red-labeled **(A)** LNCaP or **(B)** DU145 showed a significant increase in cytotoxicity over time (fold change relative to untreated control at each timepoint. *\*P*, < 0.05 by paired student’s t-test.

**Fig. S3.**
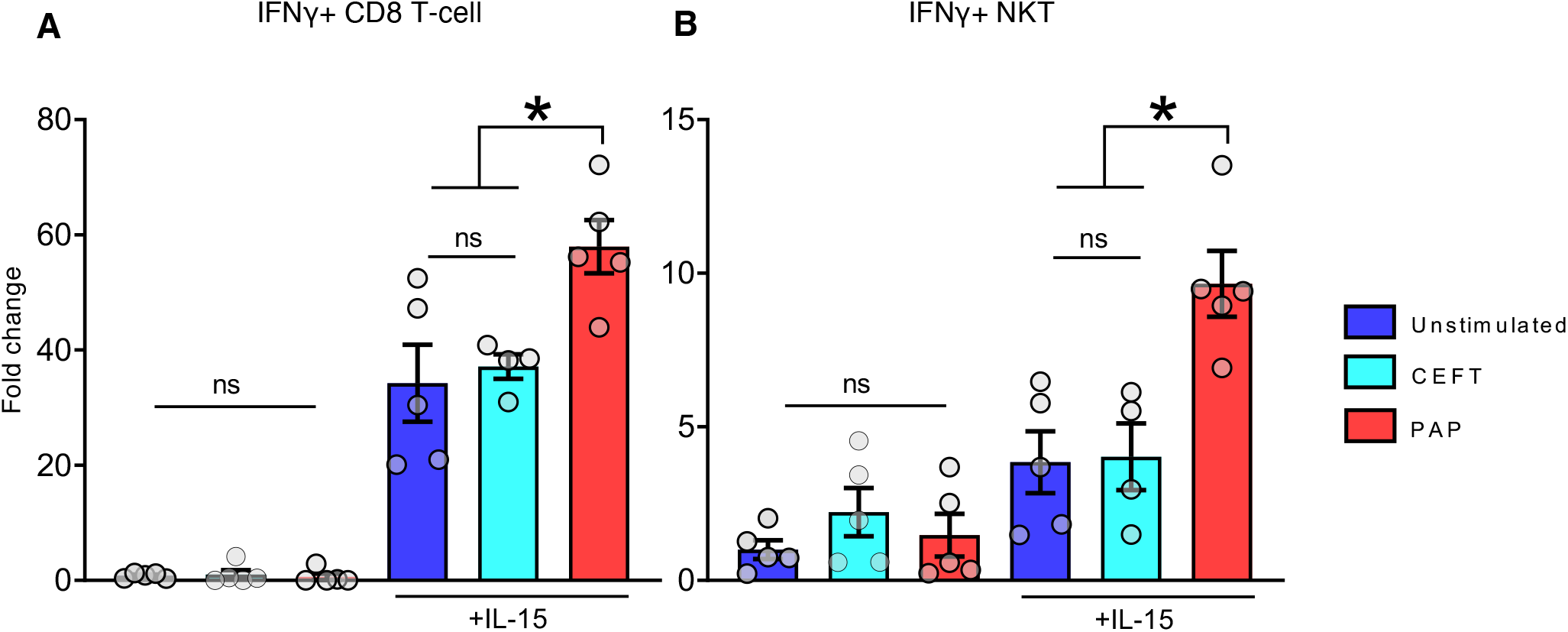
IL-15 expands antigen-specific T-cell subsets of sip-T. Sip-T from two individual mCRPC patients (n=5) were treated with prostatic acid phosphatase (PAP; 1 µg/ml) or control peptide, CEFT (1 µg/ml) alone or in combination with IL-15 (50 ng/ml). Unstimulated controls received media only. IL-15 was added on day 0 and 2 and cells harvested on day 3. Bar graphs (mean ± SEM shown) represent fold change relative to unstimulated controls. IL-15 treated sip-T showed significant expansion of IFN-γ+ **(A)** CD8 T-cells and **(B)** CD56+ NKT following PAP but not CEFT treatments in the presence of IL-15. *, *P < 0.05* by 1-way ANOVA with Tukey’s post hoc test.

**Fig. S4.**
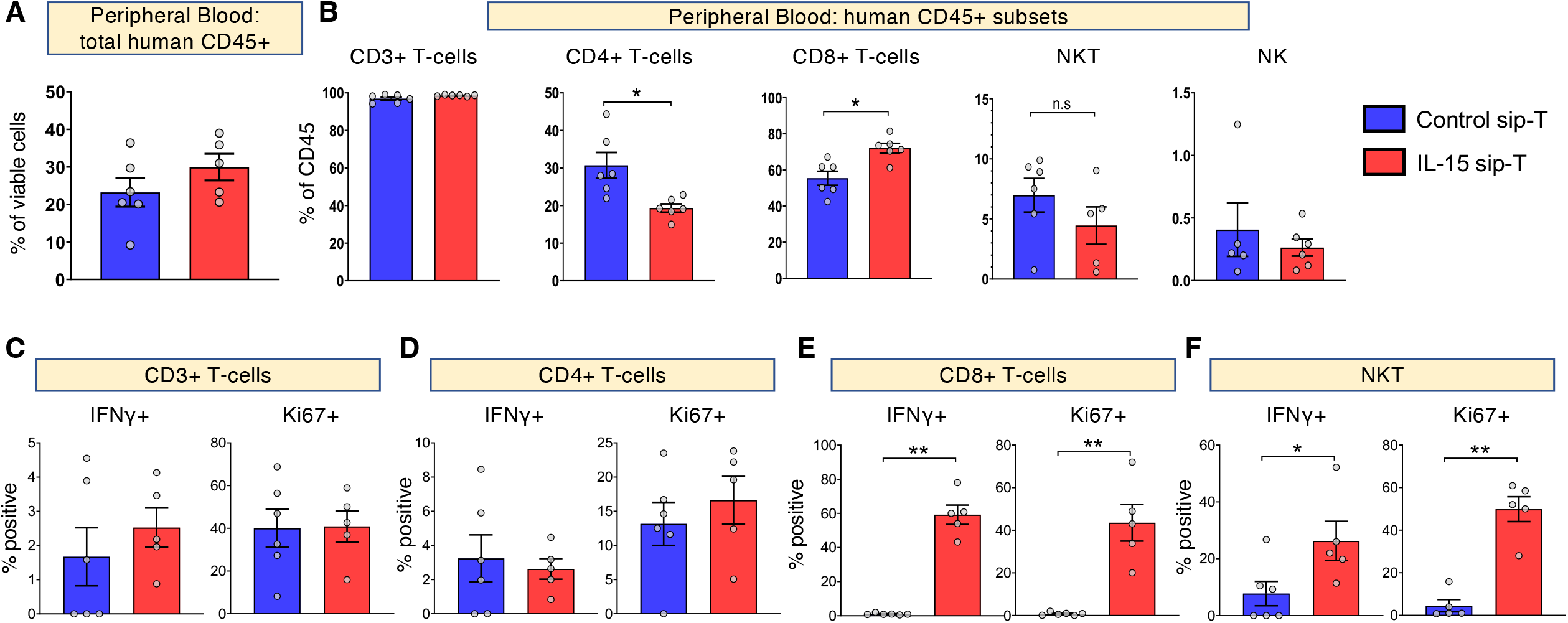
IL-15 significantly improves activation and proliferation of sip-T in peripheral blood *in vivo*. Male NOD/SCID/IL2R gamma (null; NSG) mice were implanted with 3×10^6^ LNCaP cells subcutaneously. Once average tumor size reached 50-100 mm^2^, mice received four injections of control or IL-15 treated sip-T (5×10^6^ cells intraperitoneally), seven days apart. IL-15 sip-T group also received four injections of rhIL-15 (1 µg subcutaneously) at the time of each sip-T administration. Mice were euthanized on day 28 post-initiation of sip-T treatment and peripheral blood was analyzed by flow cytometry. **(A-F)** Bar graphs of sip-T analysis in blood (mean ± SEM). Consistent with TILs data, IL-15 treated sip-T showed significantly higher activation (IFN-γ expression) and proliferation (Ki67 expression) of CD8+ T-cell and NKT subsets compared with controls. *, *P < 0.05* by Mann-Whitney.

**Table 1. Patient characteristics included in this study.**

